# Obesity causes irreversible mitochondria failure in visceral adipose tissue despite successful anti-obesogenic lifestyle-based interventions

**DOI:** 10.1101/2020.07.08.194167

**Authors:** Alba Gonzalez-Franquesa, Pau Gama-Perez, Marta Kulis, Norma Dahdah, Sonia Moreno-Gomez, Ana Latorre-Pellicer, Rebeca Fernández-Ruiz, Antoni Aguilar-Mogas, Erika Monelli, Sara Samino, Joan Miró, Gregor Oemer, Xavier Duran, Estrella Sanchez-Rebordelo, Marc Schneeberger, Merce Obach, Joel Montane, Giancarlo Castellano, Vicente Chapaprieta, Lourdes Navarro, Ignacio Prieto, Carlos Castaño, Anna Novials, Ramon Gomis, Maria Monsalve, Marc Claret, Mariona Graupera, Guadalupe Soria, Joan Vendrell, Sonia Fernandez-Veledo, Jose Antonio Enríquez, Angel Carracedo, José Carlos Perales, Rubén Nogueiras, Laura Herrero, Markus A. Keller, Oscar Yanes, Marta Sales-Pardo, Roger Guimerà, José Ignacio Martín-Subero, Pablo M. Garcia-Roves

**Affiliations:** Department of Physiological Sciences, Universitat de Barcelona, 08907 Barcelona, Spain; Novo Nordisk Foundation Center for Basic Metabolic Research, University of Copenhagen, Denmark; Biomedical Epigenomics Group, Institut d’Investigacions Biomèdiques August Pi i Sunyer (IDIBAPS), 08036 Barcelona, Spain; Grupo de Medicina Xenómica, CIBERER, Centre for Research in Molecular Medicine and Chronic Diseases (CIMUS), Universidade de Santiago de Compostela, 15782 Santiago de Compostela, Spain; Unit of Clinical Genetics and Functional Genomics, Department of Pharmacology-Physiology, School of Medicine, University of Zaragoza, CIBERER-GCV02 and ISS-Aragon, 50009 Zaragoza, Spain; Diabetes and Obesity Research Laboratory, Institut d’Investigacions Biomèdiques August Pi i Sunyer (IDIBAPS), 08036 Barcelona, Spain; Centro de Investigación Biomédica en Red Diabetes y Enfermedades Metabólicas Asociadas (CIBERDEM), 28029 Madrid, Spain; Department of Chemical Engineering, Universitat Rovira i Virgili, 43007 Tarragona, Spain; Vascular Signalling Laboratory, Program Against Cancer Therapeutic Resistance (ProCURE), Institut d’Investigació Biomèdica de Bellvitge (IDIBELL), 08907 Barcelona, Spain; Metabolomics Platform, IISPV, Department of Electronic Engineering, Universitat Rovira i Virgili, 43007 Tarragona, Spain; Institute of Human Genetics, Medical University of Innsbruck, 6020 Innsbruck, Austria; Institut d’Investigació Sanitària Pere Virgili (IISPV), 43204 Reus, Spain; Department of Physiology, CIMUS, Universidad de Santiago de Compostela, 15782 Santiago de Compostela, Spain; Laboratory of Molecular Genetics, Howard Hughes Medical Institute, The Rockefeller University, NY10065 New York, USA; Blanquerna School of Health Science, Ramon Llull University, Barcelona, Spain; Instituto de Investigaciones Biomédicas Alberto Sols (CSIC-UAM), 28029 Madrid, Spain; Neuronal Control of Metabolism Laboratory, Institut d’Investigacions Biomèdiques August Pi i Sunyer (IDIBAPS), 08036 Barcelona, Spain; Experimental 7T MRI Unit, Institut d’Investigacions Biomèdiques August Pi i Sunyer (IDIBAPS), Barcelona, Spain; Centro Nacional de Investigaciones Cardiovasculares Carlos III, Madrid, Spain; CIBERFES, 28029 Madrid, Spain; Fundación Pública Galega de Medicina Xenómica, SERGAS, Instituto de Investigación Sanitaria de Santiago de Compostela (IDIS), Santiago de Compostela, Spain; Nutrition, Metabolism and Gene therapy Group; Diabetes and Metabolism Program; Institut d’Investigació Biomèdica de Bellvitge (IDIBELL), Barcelona, Spain; Centro de Investigación Biomédica en Red Fisiopatología de la Obesidad y la Nutrición (CIBEROBN), Instituto de Salud Carlos III, 28029 Madrid, Spain; Department of Biochemistry and Physiology, School of Pharmacy and Food Sciences, Institut de Biomedicina de la Universitat de Barcelona (IBUB), Universitat de Barcelona, 08028 Barcelona Spain; Institució Catalana de Recerca i Estudis Avançats (ICREA), 08010 Barcelona, Spain; Department of Basic Clinical Practice, Universitat de Barcelona, 08036 Barcelona, Spain

**Author notes:** These authors contributed equally (Alba Gonzalez-Franquesa, Pau Gama-Perez).

**Keywords:** Obesity, metabolic plasticity, visceral adipose tissue, mitochondrial dysfunction, exercise, caloric restriction, multi-organic approach, lifestyle intervention, aging-like phenotype, metabolic fingerprint.

## Abstract

Metabolic plasticity is the ability of a biological system to adapt its metabolic phenotype to different environmental stressors. We used a whole-body and tissue-specific phenotypic, functional, metabolomic and transcriptomic approach to systematically assess metabolic plasticity in diet-induced obese mice after a combined nutritional and exercise intervention. Although most pathological features were successfully reverted, we observed a high degree of metabolic dysfunction irreversibility in visceral white adipose tissue, characterised by abnormal mitochondrial morphology and functionality. Despite two sequential therapeutic interventions and apparent global phenotypic recovery, obesity specifically triggered in visceral adipose a cascade of events progressing from mitochondrial metabolic and proteostatic defects to widespread cellular stress, which compromises its biosynthetic and recycling capacity. Our data indicate that obesity prompts a lasting metabolic fingerprint that leads to a progressive breakdown of metabolic plasticity in white adipose tissue, becoming a significant milestone in disease progression.

## Introduction

Food availability, consumption surplus and sedentarism, predominant in most parts of the modern world, are the most significant environmental challenges that western societies are facing nowadays. According to the World Health Organization (WHO), in 2016, 650 million adults were suffering from obesity, with a growing incidence that had become a significant concern in healthcare (*1*). Moreover, obesity increases the risk for most common non-communicable diseases, including neurodegenerative and cardio-vascular diseases, cancer and type 2 diabetes (T2D). Tackling the biology underlying obesity-related TD2 is a challenge, as it is a multi-organic disease, from both pathological and etiological perspectives. Surgical, pharmacological and lifestyle-based interventions partially revert the obesity phenotype, although thwarted by their failure to sustain body weight loss and health benefits and frequent relapses (*2–4*). This failure to reverse obesity suggests that cellular and metabolic plasticity, interpreted as the ability of the cells to change and efficiently adapt their phenotypes to specific environmental cues, may be impaired. This phenomenon takes place in chronic conditions and diseases as well as in ageing, but whether obesity disrupts metabolic plasticity, notwithstanding therapeutic interventions, is still a conundrum.

In this context, the LiMa (<u>Li</u>festyle <u>Ma</u>tters) project uses a comprehensive therapeutic approach focused on the systemic and tissue-specific impact of important environmental stressors (namely chronic overfeeding and physical inactivity). Our goal is to elucidate the reversibility and improvement of metabolic plasticity achieved by lifestyle interventions on the pathological changes in different organs caused by obesity and insulin resistance. Here, we present an integrative multi-tiered approach to study the impact of lifestyle interventions on obesity, and reveal that obesity triggers a progressive breakdown of metabolic plasticity in white adipose tissue which is irreversible despite apparent phenotypic reversion.

## Results

### Combined nutritional and exercise intervention reverses HFD-induced phenotype

We investigated how reversible is diet-induced obesity through lifestyle intervention. Three experimental groups were defined: control group (Ctrl); HFD-induced obese and insulin-resistant group (HFD); and a third group (Int) which, included a nutritional and exercise intervention: specifically, a calorie restriction diet substitution (with poly- and mono-unsaturated rich oils and complex carbohydrates instead of saturated fats and simple sugars) combined with treadmill training, (*see methods section for detailed group description*). HFD group showed increased body weight (Figure 1A) by expanded fat volume (Figures 1B and 1C). Also, HFD mice showed increased triglyceride content in liver and soleus skeletal muscle (Figure 1D), along with disrupted metabolic flexibility (Figures 1E and 1F) and impaired glucose metabolism (Figures 1G-H) in comparison to the control group (Ctrl group). Glucose metabolism impairment was due to insulin resistance (Figure S1A) and pancreatic dysfunction evident by increased pancreas size, impaired glucose-stimulated insulin secretion, both *in vivo* and *in vitro* (Figures 1H-L) and increased β cell mass (Figures 1K-L, Figure S1B).

**Figure 1.**
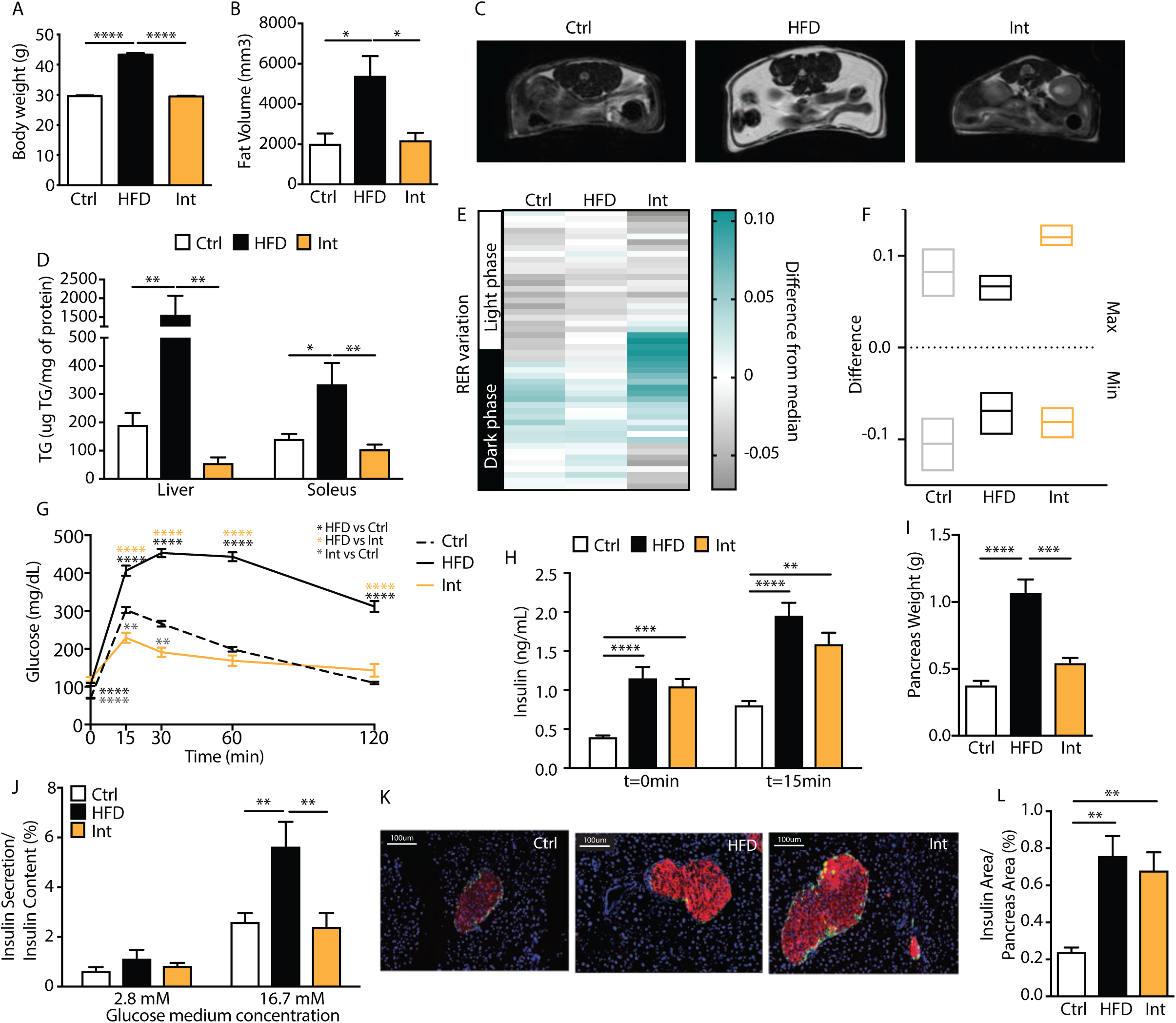
HFD- and combined intervention-induced phenotype. A: Body weight for control (Ctrl, n=80), high fat diet-fed (HFD, n=70) and intervention (Int, n=46) groups. B: Quantification of fat volume by NMR imaging and 7Teslas scan representative images per phenotype (Ctrl n=4; HFD n=4; Int n=3). C: NMR representative images for each group for quantification in (B). D: Triglycerides (TG) levels in liver: Ctrl (n=8), HFD (n=7) and Int (n= 8); and soleus: Ctrl (n= 9), HFD (n= 8) and Int (n=8). E: Group-averaged respiratory exchange ratio (RER) deviation from the 24h median for each animal for Ctrl (n=11), HFD (n=10) and Int (n=6) for 48h in metabolic cages. F: Minimum and maximal values of deviation from median (RER variation, Figure E). G: Intraperitoneal glucose tolerance test (IGTT) for Ctrl (n=54), HFD (n=48) and Int (n=12). H: Basal (t=0min) and 15min insulin levels during IGTT (Figure G) for Ctrl (n=15), HFD (n=19) and Int (n=13). I: Pancreas mean tissue weight for Ctrl (n=6), HFD (n=6) and Int (n=4). J: Glucose-stimulated (2.8 and 16.7mM) insulin secretion *in vitro* for pancreatic islets from animals in groups Ctrl (n=10), HFD (n=7) and Int (n=7). K: Immunohistochemistry representative images of pancreatic islets (nuclei-blue, insulin-red, glucagon-green) from animals in Ctrl, HFD and Int. L: Normalized pancreatic islet insulin area by pancreas section area for Ctrl (n=6), HFD (n=6) and Int (n=4). Ctrl: control; HFD: high fat diet-fed; Int: intervention. Data represented as mean ± SEM, ANOVA One-way and Post-hoc Tukey, *p<0.05, **p<0.01, ***p<0.001, and ****p<0.0001.

Combined nutritional and exercise intervention (Int group) reversed the observed pathophysiology (Figures 1A-J) except for persistent hyperinsulinemia (both fasting and after glucose stimulation) (Figure 1H), consistent with the maintained increase in β cell mass after the intervention (Figures 1K-L, Figure S1B), even though *in vitro* GSIS was fully recovered (Figure 1J). Altogether these results validated our diet-induced obesity animal model and efficiency of the combined nutritional and exercise intervention to revert obesity and partially the T2D phenotype.

### Tissue-specific pathological consequences of obesity and recovery after combined nutritional and exercise intervention

Next, we assessed the tissue-specific effect of the combined intervention on metabolic plasticity. We used a three-tiered approach to elucidate the pathophysiological impact of obesity-related insulin resistance in skeletal muscle, liver, WAT, and hypothalamus. Specifically, this approach consisted of evaluating mitochondrial respiration states (Mito); targeted expression profiles of genes related to glucose and fatty acid metabolism, mitochondrial biogenesis, function, and stress (Genes); and analysis of metabolites (Mets) (Figure 2A, Figures S2A-B and Suppl. Table 1), investigating them in the tissues mentioned above and screening for the presence or absence of reversibility.

**Figure 2.**
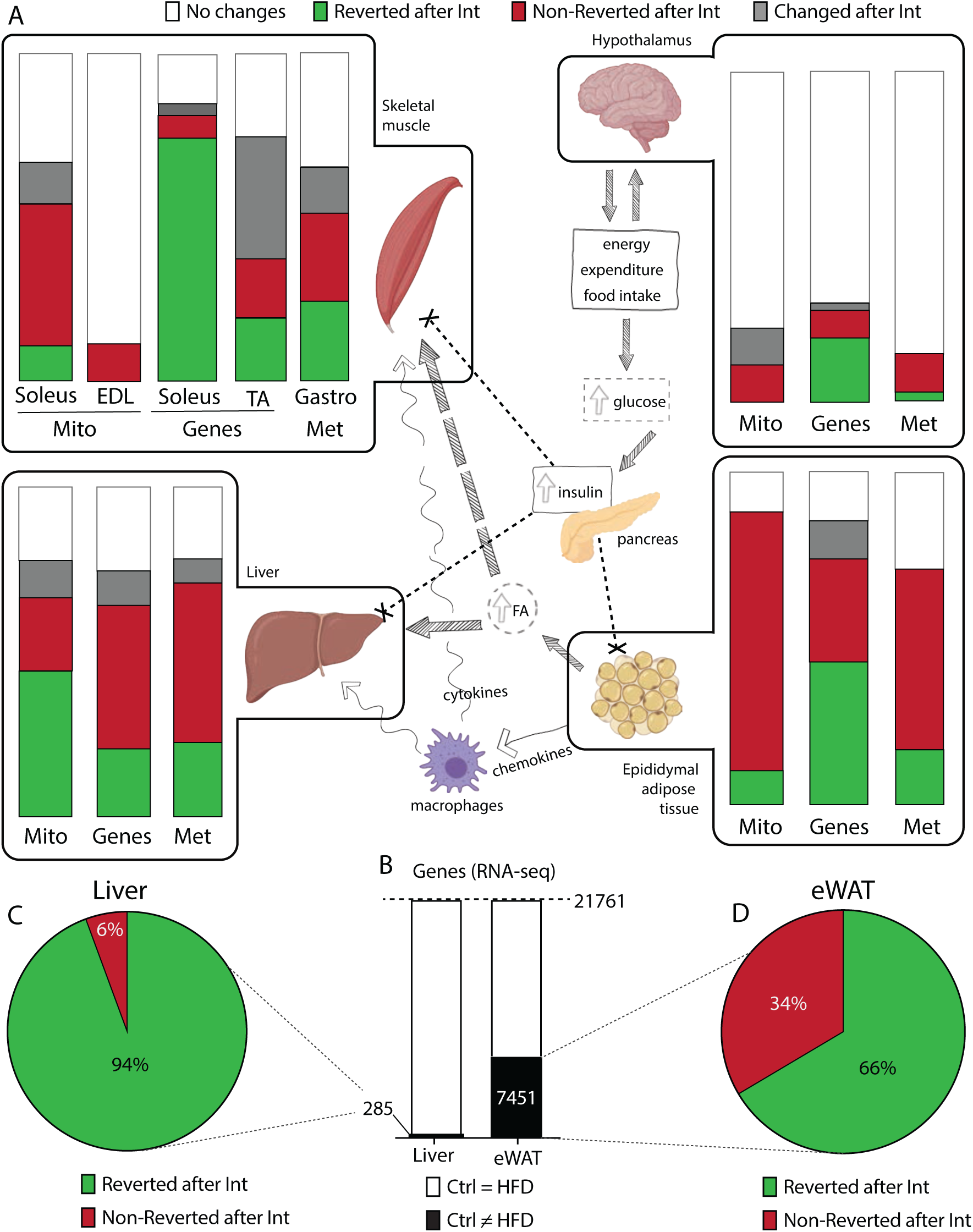
Tissue-specific effects of HFD-feeding and nutritional and exercise intervention. A: Schematic overview of the main processes taking place in tissues involved in obesity-related insulin resistance. Boxes highlight tissues analysed in this study (skeletal muscle, hypothalamus, liver and epidydimal white adipose tissue, eWAT), where we represent the changes induced by HFD and Intervention on mitochondrial respiratory states (Mito), gene expression (Genes) and metabolites (Met). Each bar corresponds to the 100% of respiratory states/genes/metabolites analysed. Bands of different colors represent fractions of respiratory states/genes/metabolites showing different patterns of change: white – no changed; green – changed after HFD-feeding and reverted with intervention; red – changed after HFD-feeding and not reverted with intervention; grey – changed uniquely with intervention. B: Bar graph for RNA-seq in liver and eWAT. The black area represents gene transcripts with changed expression after HFD-feeding. C-D: Percentage of gene transcripts reverted (green) or non-reverted (red) in liver (C) and eWAT (D) after the combined intervention.

Limited changes were observed in hypothalamus regardless of HFD-feeding or intervention status. The intervention induced significant changes in glycolytic skeletal muscle (tibialis anterior), consistent with the switch to an oxidative phenotype promoted by the implementation of the exercise program. Mice on HFD-feeding induced substantial metabolic changes (combined green and red bars) in oxidative skeletal muscle (soleus). Of note, most of these metabolic changes were reverted across the different tiers in muscle after the intervention (green bars). Liver and epidydimal WAT (eWAT) however, suffered profound metabolic changes upon HFD-feeding (combined green and red bars) and even though the whole body improved metabolic phenotype, these two tissues specifically exhibited sparse or non-reversion (red bars) in all three phenotypical tiers (Figure 2A). To further assess the irreversible loss of metabolic plasticity, we performed next-generation RNA sequencing (RNA-seq) in liver and eWAT before and after the intervention. Differential expression analysis revealed that, upon HFD-feeding, eWAT transcriptome was much more perturbed than the liver transcriptome (7,451 vs 285 genes, respectively) (Figure 2B) and that upon intervention, the eWAT transcriptome exhibited a lesser degree of reversion (66% as compared to 94% in the liver) (Figures 2C and 2D).

Collectively, these results argued for a tissue-specific heterogeneous susceptibility to obesity-induced defects in metabolic plasticity, and that eWAT specifically showed the highest degree of irreversibility after a combined nutritional and exercise intervention.

### Incomplete remodeling of eWAT after the combined intervention

To further characterise the changes accounting for obesity-related metabolic irreversibility in eWAT, we studied the transcriptome. We identified downregulated (Figure 3A) and upregulated genes (Figure 3G) after HFD-feeding. Within the 3,869 downregulated genes, 1,036 recovered, 1,464 partially recovered, and 1,369 remained downregulated after the intervention (Figure 3A). Gene ontology enrichment analysis showed that non-reverted transcripts were related to biological processes such as biosynthesis, metabolism, and mitochondrial morphogenesis, organisation and transport (Figure 3B). In line with this, enrichment for cellular components identified genes involved in mitochondrial inner membrane (Figure S3A). Mitochondrial inner membrane is structured in *cristae*, whose density was significantly reduced after HFD-feeding and, unexpectedly, further decreased after the intervention (Figure 3C-D, Figure S3B). Decrease in cristae density was observed irrespective of maintenance of mitochondrial aspect ratio (Figure S3C). Additionally, processing of OPA1, a protein located in the inner mitochondrial membrane and master regulator of cristae architecture, revealed the same irreversible decrease after obesity and the intervention (Figure 3E). High-resolution respirometry studies in permeabilised eWAT confirmed the functional relevance of defects in cristae density revealing decreased respiratory capacity (Figure 3F).

**Figure 3.**
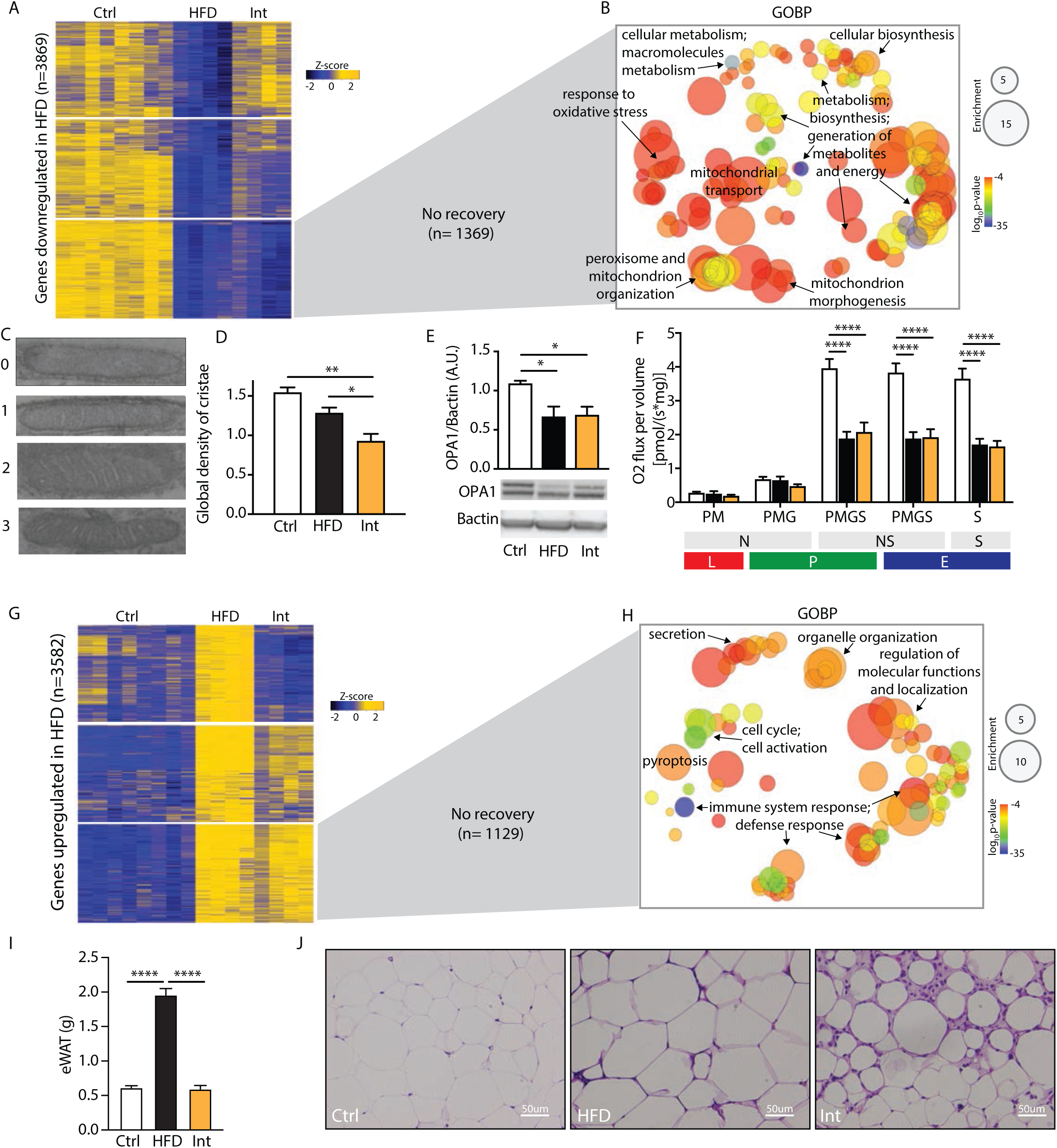
Impairment of adipose tissue plasticity. A: Heatmap representing z-scores for RNA-seq cluster of transcripts downregulated after HFD-feeding (Ctrl n=8, HFD n=4, Int n=4). B: Gene ontology biological process (GOBP) enrichment analysis for the 1369 transcripts downregulated in HFD and Int. C: Electronic microscopy representative images of mitochondria cristae frequency, category 0, 1, 2 or 3 (see Methods section for category definition). D: Cristae density score according to categories detailed in C (Ctrl n=3, HFD n=4, Int n=4). E: Normalised protein expression of OPA1 (Ctrl n=7, HFD n=9, Int n=9). F: Mitochondrial respiratory states for eWAT homogenate (Ctrl n=13, HFD n=9, Int n=10). Substrates; P: pyruvate, M: malate, G: glutamate and S: succinate. Pathways; N: NADH electron transfer-pathway, S: Succinate dehydrogenase-linked pathway and NS: NS-linked pathways. Respiratory rates; L: Leak-state; P: Oxidative phosphorylation (OXHPOS)-state; E: Electron transfer (ETS)-state (*35*). G: Heatmap representing z-scores for RNA-seq cluster of transcripts upregulated after HFD-feeding (Ctrl n=8, HFD n=4, Int n=4). H: Gene ontology biological process (GOBP) enrichment analysis for the 1129 transcripts upregulated in HFD and Int. I: Epididymal WAT weight for control (Ctrl n=32, HFD n=32, Int n=26). J: Representative images for hematoxylin and eosin eWAT staining for each experimental group (Ctrl n=3, HFD n=4, Int n=4). Ctrl: control; HFD: high fat diet-fed; Int: intervention. Data represented as mean ± SEM, ANOVA One-way and Post-hoc Tukey, *p<0.05, **p<0.01, ***p<0.001, and ****p<0.0001.

From the 3,582 differentially upregulated genes after HFD-feeding (Figure 3G), 1,064 achieved a complete reversion, 1,389 were partially reverted and 1,129 transcripts did not change after the intervention. Gene ontology enrichment analysis of these non-reverted transcripts revealed biological processes associated with the immune system and defense response (Figure 3H). To further validate these findings, we studied tissue immune infiltration in eWAT. The histological analysis revealed hypertrophied adipocytes surrounded by crown-like structures (composed by immune cells) in the HFD group (Figure 3J). Lastly, despite the improved systemic metabolic phenotype and the reduction of adipose tissue weight promoted by the intervention (Figure 3I), the histology of eWAT in these animals revealed a persistent alteration with the presence of dead adipocytes and massive immune cell infiltration (Figure 3J). These findings were corroborated by an increase in Caspase3- and Mac2-positive cells (Figure 4D, middle and lower panel, Figure S4C), markers of apoptosis and macrophages, respectively. Altogether, these results suggested an active eWAT remodeling after the intervention, potentially explaining the observed metabolic derangements.

**Figure 4.**
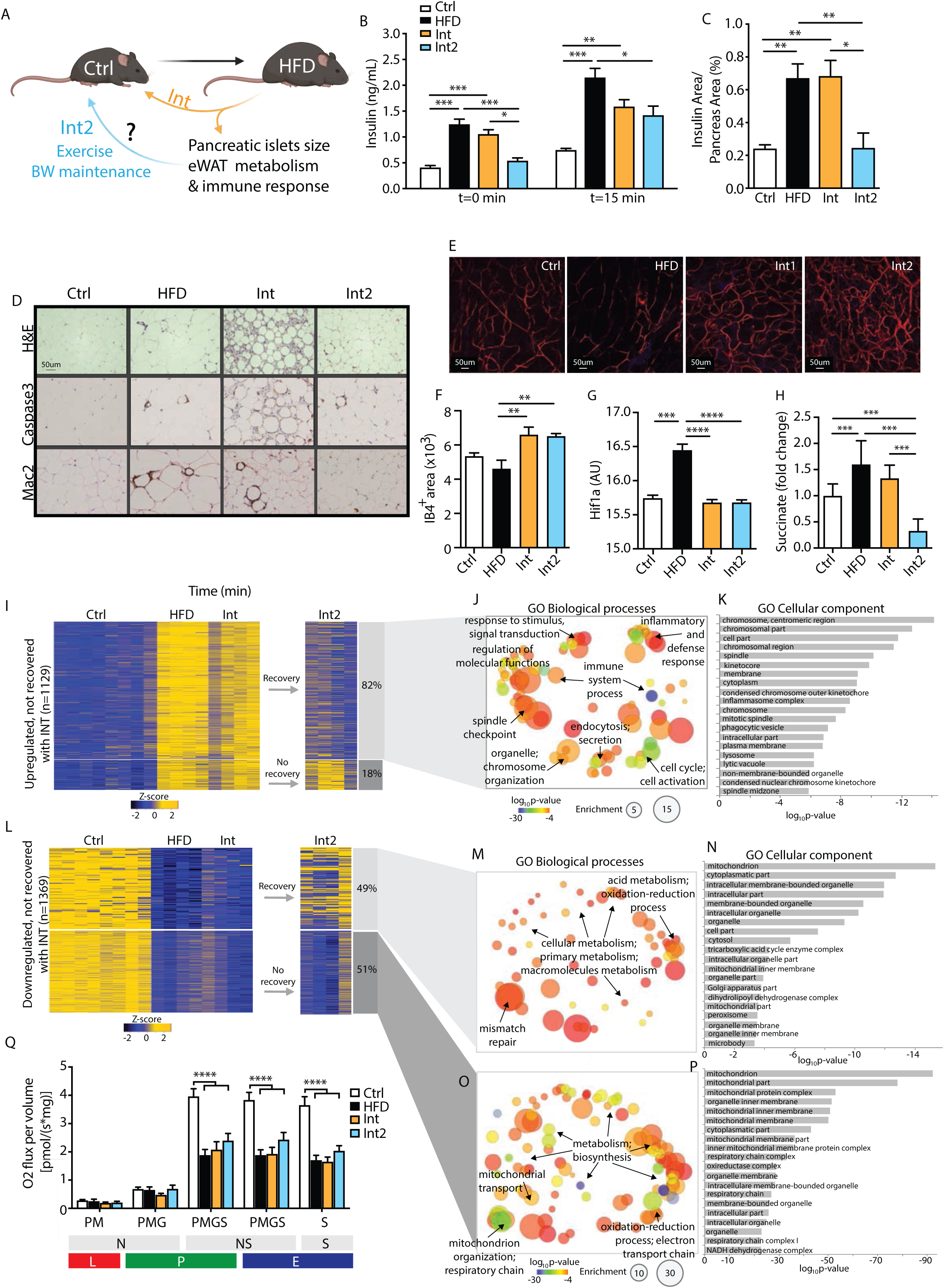
Mitochondrial deterioration in remodeled white adipose tissue. A: Scheme with the 4 experimental groups indicating the non-reverted parameters after the first intervention (Int) and the rationale to then include a second intervention (Int2). B: Basal (t=0min) and 15min insulin levels during IGTT (Ctrl n=28, HFD n=30, Int n=14, Int2 n=10). C: Normalised pancreatic islet insulin area by total pancreas area for Ctrl n=6, HFD n=6, Int n=4, Int2 n=4. D: Representative images for hematoxylin and eosin eWAT, Caspase3- and Mac2-positive cells staining in eWAT for each experimental group (Ctrl n=3-4, HFD n=4, Int n=3-4, Int2 n=4-5). E: Representative of IB4-positive stained blood vessels in flat-mounted eWAT depots from each experimental group. F: IB4-positive area quantification (Ctrl n=4, HFD n=3, Int n=4, Int2 n=6). G: Normalized Hif1-α gene expression in eWAT (Ctrl n=8, HFD n=4, Int n=4, Int2 n=4). H: Succinate metabolite levels in eWAT (Ctrl n=15, HFD n=13, Int n=6, Int2 n=6). I: Heatmap representing z-scores for RNA-seq cluster of transcripts upregulated after HFD-feeding and not recovered after Int (Ctrl n=8, HFD n=4, Int n=4, Int2 n=4). J and K: Gene ontology biological processes (J) and cellular components (K) enrichment of the genes clustered in 4I that revert after Int2. L: Z-score for RNA-seq cluster of transcripts downregulated after HFD-feeding and not recovered after Int (Ctrl n=8, HFD n=4, Int n=4, Int2 n=4). M and N: Gene ontology biological processes (M) and cellular components (N) enrichment of the genes clustered in 4L that revert after Int2. O and P: Gene ontology biological processes (O) and cellular components (P) enrichment of the genes clustered in 4L that do not revert after Int2. Q: Mitochondrial respiratory states for eWAT homogenate (Ctrl n=13, HFD n=9, Int n=10, Int2 n=13). Substrates; P: pyruvate, M: malate, G: glutamate and S: succinate. Pathways; N: NADH electron transfer-pathway, S: Succinate dehydrogenase-linked pathway and NS: NS-linked pathways. Respiratory rates; L: Leak-state; P: Oxidative phosphorylation (OXHPOS)-state; E: Electron transfer (ETS)-state. Ctrl: control; HFD: high fat diet-fed; Int: intervention; Int2: second intervention. Data represented as mean ± SEM, ANOVA One-way and Post-hoc Tukey, *p<0.05, **p<0.01, ***p<0.001, and ****p<0.0001.

### Mitochondrial dysregulation persists in morphologically remodeled eWAT

The negative energy balance induced by exercise and nutritional intervention allowed for a global metabolic phenotypical recovery except for the increased pancreatic islets size (Figures 1K and 4A) and eWAT mitochondria dysfunction and immune cells infiltration (Figures 3 and 4A). After the partial achievements of sustained weight loss and improved physical fitness, we hypothesised that it would be necessary an extended milder intervention to recover the eWAT and pancreatic islets completely. Therefore, in a separate group of mice, we conducted an extra five weeks intervention following the same diet but undergoing pair-feeding and a less intense training program aiming at bodyweight maintenance (Figure 4A).

After this second phase of the intervention (Int2 group) (Figure S4A) the body weight was maintained, and glucose tolerance was comparable to the Int group (Figure S4B). Furthermore, the second intervention successfully decreased fasting insulin and glucose-stimulated insulin levels (Figure 4B) in line with reduced β cell mass (Figure 4C). The second intervention also improved eWAT atrophy and immune cell infiltration (Figure 4D, upper panel), along with a progressive reduction of Caspase3- and Mac2-positive cells (Figure 4D, middle and lower panel, and Figure S4C).

Adipose tissue expansion negatively influences tissue remodeling and functionality by limiting oxygen and nutrient supply to adipocytes. Oxygen limitation triggers an increase in succinate levels, which stabilises and activates HIF1-α, a master regulator of the cellular response to hypoxia and angiogenesis. Thus, we next examined whether the intervention had remodeled vascularisation parameters in eWAT. Unexpectedly, confocal microscopy studies revealed enhanced eWAT vascularisation after both interventions (Figures 4E and 4F). Also, analysis of critical mediators of angiogenesis in the Int2 group, revealed normalisation of *Hif1α* gene expression (Figure 4G), *Vegfα* and *Vegfr* gene expression (Figure S4D and S4E), and reduced levels of succinate (Figure 4H).

Additionally, we evaluated whether this eWAT morphological recovery after the second intervention was accompanied by reversion of gene expression patterns by RNA-seq. Specifically, out of the 1,129 transcripts upregulated after HFD and Int (Figure 3G), 82% had reverted to Ctrl values after Int2 (Figure 4I). Gene ontology analysis of these genes that recovered their expression revealed enrichment in biological processes linked to organelle and cytoskeleton organisation, immune system, and inflammatory and defense response (Figure 4J and 4K). Besides that, from the 1,369 transcripts downregulated after HFD and Int (Figure 3A), 49% were recovered after Int2 while 51% still remained downregulated (Figure 4l). Gene ontology of the set of genes that were unable to restore their levels, showed, again, a significant enrichment in mitochondrial-related biological processes, such as electron transport chain, energy metabolism, biosynthesis, and mitochondrial organisation and transport (Figures 4O and 4P). Consistent with this result, mitochondrial respiratory capacity remained reduced after the second intervention (Figure 4Q). Altogether, these results demonstrate that metabolic and mitochondrial derangements in eWAT were resilient, and persisted even after resolution of morphological remodeling of the tissue, arguing for defects in mitochondria to be the main determinants of metabolic plasticity breakdown in eWAT.

### Enduring mitochondrial derangements as a driver of WAT metabolic plasticity breakdown

Next, we speculated that additional transcriptional changes cascading downstream the HFD-induced phenotype during the interventions might help understand the observed metabolic scenario.

Three clusters were clearly defined amongst the genes irreversibly downregulated across the experimental groups: i) 893 genes irreversibly decreased with HFD-feeding (Figure 5A, Cluster A), ii) 601 genes irreversibly decreased from Int (Cluster B) and, iii) 678 genes with partially reduced expression in Int that became significantly downregulated only after Int2 (Cluster C). Gene ontology revealed biological processes and cellular components enriched in these three clusters (Figures 5B-C). Genes irreversibly downregulated since the onset of diet-induced obesity and insulin resistance (Cluster A) related to metabolic and biosynthetic processes at the mitochondria and peroxisome level. Additionally, several other biological processes enriched in Cluster A, such as lipid metabolism, adipocyte differentiation or apoptosis, were further amplified in gene number size after the interventions (Clusters B and C, inner circles). Genes downregulated after the first intervention (Cluster B) were also related to additional processes such as RNA and glucose metabolism; protein folding and transport to Golgi and mitochondria, as well as mitochondrial organisation. Of significant relevance is the fact that previously downregulated genes were still affected or further amplified even after the second intervention (Cluster C). Also, genes from Cluster C showed new significant enrichments, such as in mitochondrial organisation, proteasome complex and ribosomal biogenesis. Hence, these findings indicate the irreversible course towards failure of transcriptional changes even after reversal of the diet-induced obese and insulin resistant phenotype during the sequential interventions. These changes lead to a progressive deterioration of eWAT functionality and pointed to the mitochondria as the primary organelle affected, triggering further transcriptional changes of genes involved in Golgi apparatus and Endoplasmic Reticulum (Figure 5C).

**Figure 5.**
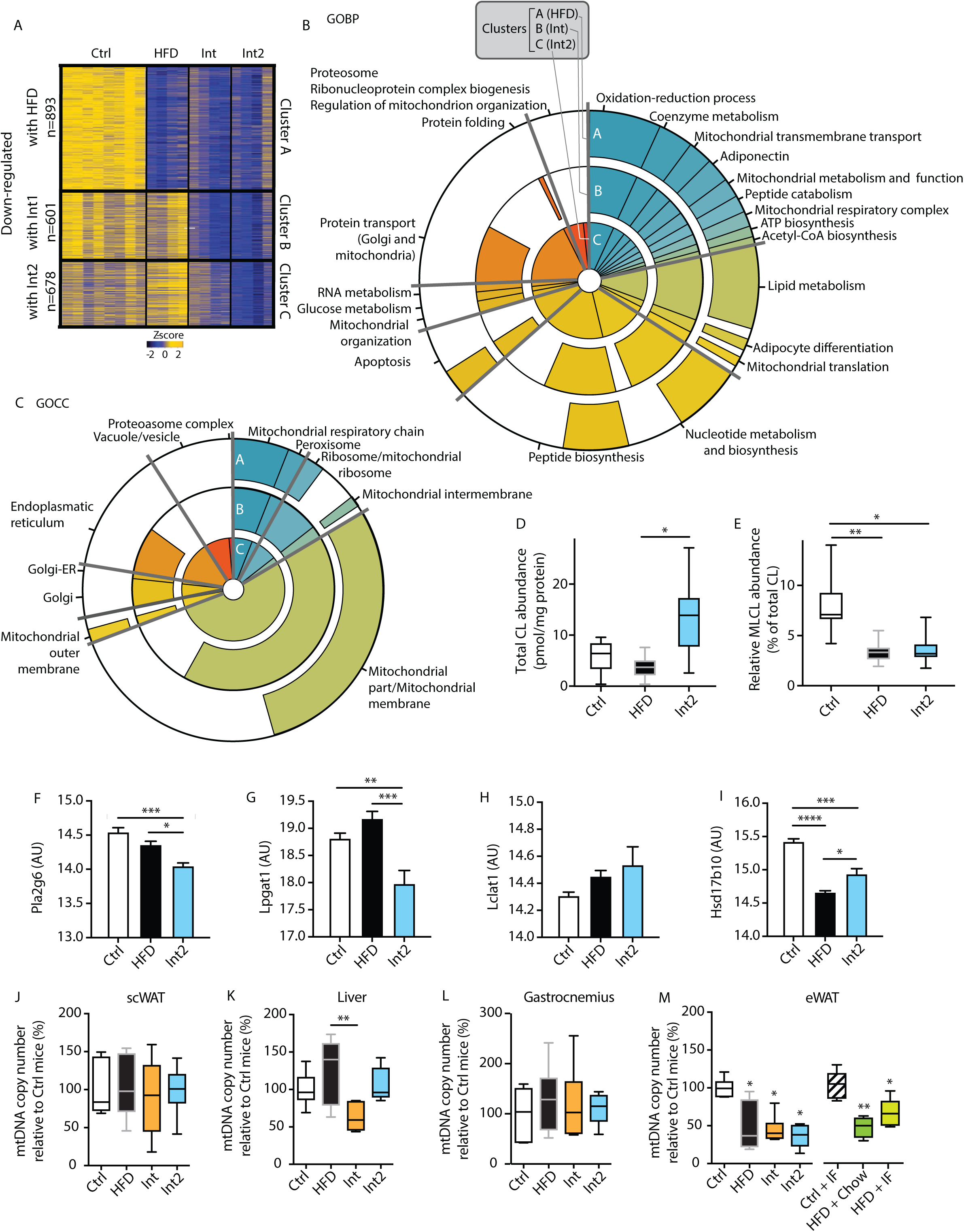
Metabolic plasticity breakdown in eWAT. A: Z-score for RNA-seq clusters of transcripts sequentially downregulated after HFD-feeding or the interventions, namely clusters A, B and C (Ctrl n=8, HFD n=4, Int n=4, Int2 n=4). B and C: Radar chart showing the gene ontology biological processes (B) and cellular components (C) enrichment after HFD-feeding (cluster A) or the interventions (clusters B and C). Pie proportion indicates gene size for each term or terms-groups indicated in the outside of the radar chart. Different colors represent different terms or terms-groups. D and E: Total cardiolipin (CL) abundance (D) and Monolysocardiolipin (MLCL) relative abundance (E) (Ctrl n=5, HFD n=10, Int2 n=8). F-I: Normalized gene expression of Pla2g6 (F), Lpgat1 (G), Lclat1 (H) and Hsd17b10 (I) in eWAT (Ctrl n=8, HFD n=4, Int n=4, Int2 n=4). J-M: Mitochondrial DNA (mtDNA) copy number in subcutaneous white adipose tissue (J), liver (K), skeletal muscle gastrocnemius (L) and epididymal WAT (M) (Ctrl n=6, HFD n=6, Int n=6, Int2 n=5, Ctrl+IF n=8, HFD+Chow n=4, HFD+IF n=10, * indicate significant differences with Ctrl group); *normalized by Ctrl mice* refers to eWAT Ctrl group (M). Ctrl: control; HFD: high fat diet-fed; Int: intervention; Int2: second intervention; Ctrl+IF: control + dietary intervention; HFD+Chow: high fat diet-fed + chow Ad libitum; HFD+IF: high fat diet-fed + dietary intervention. Data represented as mean ± SEM, ANOVA One-way and Post-hoc Tukey, *p<0.05, **p<0.01, ***p<0.001, and ****p<0.0001.

In agreement with the alterations in the inner mitochondrial membrane, cardiolipidomic analysis revealed that both HFD and Int2 group had decreased monolysocardiolipin (MLCL) relative abundance, even when total cardiolipin (CL) levels were already recovered (Figures 5D and 5E). Low MLCL levels likely indicate decreased cardiolipin remodeling rates as reported by CL deacylation-reacylation cycle genes being downregulated (i.e., *Pla2g6*, *Hsd17b10* and *Lpgat1*) or slightly increased (i.e., *Lclat1*) (Figures 5F-I). Also, a substantial reduction in mitochondrial DNA (mtDNA) copy number in eWAT (Figure 5M) supported previous findings and strengthened mitochondrial derangements as a critical component in WAT metabolic plasticity breakdown. Of note, other adipose depots such as subcutaneous WAT (scWAT, Figure 5J), liver (Figure 5K) and skeletal muscle gastrocnemius showed no changes in mtDNA (Figure 5L).

Finally, to identify the potential trigger of mtDNA reduction, we evaluated additional experimental groups in different dietary regimes aimed to overcome obesity. Intriguingly, all animals subjected to HFD-feeding also presented a decrease in mtDNA copy number regardless of the downstream intervention (namely intermittent fasting or switch to chow diet) (Figure 5M). On the contrary, such reduction was not observed in animals that were not previously obese but followed the same dietary regime (Figure 5M), suggesting that the stress induced by HFD could trigger a permanent damage. Collectively, our results reveal that phenotypically healthy mice after reversal of obesity show tissue-specific breakdown of metabolic plasticity, and highlight eWAT as the primary target. This breakdown is primarily characterised by a sequential and unresolved mitochondrial deterioration, further aggravating other cellular functions even after combined exercise and nutritional interventions.

## Discussion

Obesity-related T2D is a chronic and degenerative disease that involves the interplay amongst and failure of several organs and tissues at both its onset and progression. This level of complexity requires a multidisciplinary and integrative experimental approach to capture interorgan communication and elucidate the vulnerability and resilience of organs leading to a global metabolic failure (*5, 6*). Thus, the current study uses a multidisciplinary and multi-organ approach to analyse an experimental combined intervention that proved to be successful in recovering nearly all phenotypical metabolic alterations in a diet-induced obese animal model. This animal model resembles the early stages of human obesity-related T2D pathology (*7, 8*). Moreover, this study provides valuable information in that it shows that lifestyle interventions are powerful treatments to overcome obesity and associated comorbidities. In addition, it allows for assessment of not only the phenotypic impact of HFD, but also how its metabolic fingerprint affects different tissues.

Our approach identifies the visceral WAT as the most affected tissue with metabolic derangements in obesity defined by its irreversible loss of functional plasticity despite apparently successful therapeutic interventions. An in-depth analysis of the global transcriptome in eWAT from obese mice reveals a generalised downregulation of metabolic processes mainly affecting mitochondria, and particularly its inner membrane and oxidative phosphorylation system (*9, 10*). The negative impact of chronic overfeeding in WAT mitochondria and how defective mitochondria performance influence WAT functionality has been extensively studied (*11–13*). The critical observation and novel finding here is that a combined nutritional and exercise intervention, powerful to reverse obesity systemically and improve tissue-specifically into a healthy phenotype, leads to a long-lasting obesity-related mitochondrial deterioration in visceral WAT that becomes irreversible.

To elucidate whether this deterioration was due to an intrinsic mitochondrial defect or a consequence of a loss of tissue plasticity, we studied additional functional parameters. A considerable amount of evidence supports the requirement of an optimised degree of WAT expandability to preserve whole body homeostasis when faced with energy surplus challenges (*14*). Factors such as early angiogenesis and inflammation in response to mild tissue hypoxia are critical requirements for a healthy WAT expansion (*15, 16*). However, the chronicity of metabolic overload stress promoted by extended overfeeding periods can compromise tissue plasticity, creating a maladaptive response characterised by persistent oxidative damage, inadequate angiogenic response, severe hypoxia, adipocyte death, immune infiltration and fibrosis (*17*). Although these pathogenic events have been described, the mechanisms involved in the remodeling during the shrinkage promoted by weight loss remains controversial (*18–23*). On this respect, we observed evidence of hypoxia (namely accumulation of succinate levels and *Hif1-α* overexpression) in WAT of obese mice, without a parallel VEGF-mediated proper vascularisation, as well as marked dystrophy-like tissue architecture after weight loss. This suggests an evident defect in the remodeling mechanisms of adipose tissue. By implementing the second phase of intervention with maintenance of low body weight, all these parameters were normalised, thus indicating the completion of the remodeling process. Nonetheless, despite such improvements in tissue architecture and vascularisation, mitochondrial defects were still present both at a transcriptional and functional level. Altogether, these results reinforce the novel finding of a permanent intrinsic mitochondrial damage in eWAT regardless of the tissue remodeling capacity (tissue plasticity).

The magnitude and specificity of the mtDNA reduction observed in eWAT amongst different experimental groups and tissues is a robust indicative sign of severe mitochondrial disturbances at the tissue specific-level and points at HFD-feeding as the driving force for such mitochondrial deterioration. Identification of the causative mechanisms leading to this irreversible structural, functional and transcriptional mitochondrial damage is challenging, due to the multifactorial components occurring under physiological conditions. Nonetheless, our findings reveal a series of events leading to mitochondrial damage and impaired tissue metabolic plasticity. It is well established that proper fine-tuning between mitochondrial biogenesis and recycling mediated by quality-control mechanisms is essential to ensure optimal functionality (*24*). However, critical perturbations in mitochondrial quality control could potentially cause cellular dysfunction involving protein import deficiencies and leading to cellular proteostatic defects and an ageing phenotype (*25*). In line with this, a recent paper by Yu *et al* demonstrated how, in the context of obesity, a disrupted protein import machinery in adipocytes is related to mitochondrial malfunction (*26*). Accordingly, the in-depth Gene Ontology analysis performed at different time-points shows that the downregulation of metabolic processes, initially confined to mitochondria, is spread to biological processes related to protein transport and folding, affecting other cellular compartments such as Golgi, Endoplasmic Reticulum and the ubiquitin and proteasome system. Overall, these data are indicative of the spreading of original mitochondrial stress to other structures and processes, worsening global cellular stress and the ability to degrade and recycle damaged proteins or organelles. This finding is especially relevant in the context of a healthy phenotype at a systemic level after the intervention, in which HFD seems to trigger irreversible oxidative damage in eWAT that ultimately progresses and causes the breakdown of metabolic plasticity.

A reduced capacity to degrade or recycle cellular components could answer one of the most interesting questions arising from our results: why are such disrupted mitochondria still present in this tissue and not targeted for recycling through mitophagy processes? The observed defective cardiolipin remodeling adds insight in that direction. Indeed, cardiolipin remodeling plays a critical role in the link between oxidative stress (e.g. ROS) and mitochondrial dysfunction in different metabolic tissues (*27*). Under physiological conditions, this remodeling process links mitochondrial cardiolipin architecture to the phospholipid environment of the cell by a tissue-independent specificity mechanism that is not primarily controlled via gene expression (*28*). However, this holds as long as none of the essential components for cardiolipin remodeling becomes rate-limiting. In this respect, lysocardiolipin acyltransferase 1 (LCLAT1) has been suggested as a pivotal enzyme involved in the pathological remodeling of cardiolipin in various age-related pathological conditions including diet-induced obesity and T2D. Thus, LCLAT1 expression tendency to increase aggravates the vicious cycle among mitochondrial dysfunction, ROS damage, and defects in mitochondrial recycling capacity (*29–31*). Therefore, the combination of different data sets not only evidences a deficiency in cristae formation but also suggests a defective clearance of dysfunctional mitochondria as a plausible explanation for this loss of metabolic plasticity.

In line with our findings, Hahn *et al* recently reported that, before any other tissue, WAT of mice fed ad libitum exhibited significant age-related effects at a transcriptional level (*32*). The study also identified a WAT-specific nutritional memory that impeded its metabolic reprogramming when switched to calorie restriction. Although this study was focused in late-life calorie restriction and longevity in mice, it revealed an ageing phenomenon that shares similarities with the disease-oriented approach in our research (namely diet-induced obesity). Altogether, we hypothesise that the stress promoted by HFD might hasten adipocytes to enter in a premature ageing-like state triggered by subsequent mitochondrial dysfunction, unsuccessful mitoprotein-induced stress response and compromised cellular renewal capacity. Overall, all these events agree with age-related processes reported in other chronic degenerative diseases (*33, 34*).

To summarise, we demonstrate that nutritional and exercise interventions, resembling a healthy lifestyle in humans, can revert a diet-induced obese and insulin resistant phenotype in mice. However, the broad spectrum of data layers, studied tissues and phenotypical characterisation performed in this study indicates a hierarchy in the way organs are vulnerable and resilient. In this sense, it reveals a metabolic plasticity breakdown in white adipose tissue. Thus, mitochondrial deterioration in visceral WAT suggests an initial and critical point of no return in metabolic disease progression, and therefore a relevant target for prevention and early treatment. In the context of obesity, trigger of these processes could be therefore avoided by implementing and advocating for early lifestyle interventions in individuals at risk.

## Acknowledgements

This work has been supported by Ministerio de Ciencia e Innovación (MICINN) Grant BFU2011-24679 (P.M.G.-R.); Instituto de Salud Carlos III (ISCIII) Grant PI15/00701 (P.M.G.-R.) cofinanced by the European Regional Development Fund ‘‘A way to build Europe’’; Government of Catalonia Suport Grups de Recerca AGAUR 2017-SGR-204 (to J.C.P. and P.M.G-R.), 2017-SGR-736 (to J.I.M.-S.) and 2017-SGR-896 (to A.A.-M., M.S.-P., and R.G.). M.M. was recipient of a MINECO grant RTI2018-093864-B-I00. M.A.K. was funded by a Medical University of Innsbruck Start Grant. M.C. was funded by the European Research Council (ERC) under the European Union’s Horizon 2020 research and innovation programme (grant agreement 725004). L. H. was funded by MINECO grants SAF2013-45887-R, SAF2017-83813-C3-1-R (granted to Dolors Serra (DS) and L.H.), co-funded by the European Regional Development Fund, the Centro de Investigación Biomédica en Red de Fisiopatología de la Obesidad y la Nutrición (CIBEROBN) (Grant CB06/03/0001 to DS), the Government of Catalonia (2017SGR278 to DS), the Fundació La Marató de TV3 (201627-30 to DS), the European Foundation for the Study of Diabetes (EFSD)/Janssen-Rising Star and L’Oréal-UNESCO “For Women in Science” research fellowships. P.M.G.-R. was a recipient of a Ramon y Cajal contract (RYC-2009-05158) from MICINN. A.G-F. was a recipient of “Beca de Formació de Personal Investigador de l’IDIBAPS” fellowship and she is currently supported by an unconditional donation from the Novo Nordisk Foundation (NNF) to NNF Center for Basic Metabolic Research (NNF18CC0034900). P.G-P. was a recipient of predoctoral fellowship from the Universitat de Barcelona (APIF-UB). E.S-R. was recipient of a predoctoral fellowship from Ministerio de Economia y Empresa (MINECO) grant BES-2013-062796. This work was partially developed at the Esther Koplowitz Centre (CEK, Barcelona, Spain).

## Materials and Methods

### Animals

C57BL/6JOlaHSD male mice purchased from Envigo (Indiana, IN, USA) were kept on a 12 h light/12 h dark cycle. Mice were assigned to two groups fed with different diets for 16weeks: control group (*Ctrl*; fed with chow diet Teklad Global 14% Protein Rodent Maintenance Diet by Envigo; final inclusion criteria: body weight (BW) < 33 g, fasting normoglycemia, fasting normal insulin, glucose tolerance and insulin sensitivity); and a high-fat diet-fed group (*HFD*; D12451 by Research Diets, New Brunswick, NJ, USA), final inclusion criteria: BW> 37 g, fasting hyperglycemia, fasting hyperinsulinemia, glucose intolerance and insulin resistance). After 16 weeks on HFD, a subgroup of *HFD* mice was assigned to the intervention group (*Int*, fed with Preliminary Formula Rodent Diet with 45 kcal% Fat and Modification with Flaxseed and Olive Oil by Research Diets, New Brunswick, NJ, USA); and a further intervention was performed (*Int2*, fed with the same diet as the *Int* group), see intervention details below. All animal procedures were approved by the local ethics committee, Comitè Ètic d’Experimentació Animal at Universitat de Barcelona and the Departament d’Agricultura, Ramaderia, Pesca, Alimentació i Medi Natural at the Generalitat de Catalunya; complying with current Spanish and European legislation.

### Nutritional and exercise intervention

Nutritional and exercise intervention (*Int* group) was initially for 5 weeks. The daily energy intake was different during the first week compare to resting weeks, being mice fed with 80% and then 100% in kcal of the daily energy intake determined for a *Ctrl* animal, respectively. Intervention diet aimed at maintain energy content from fat (45%) and included several nutritional modifications: replacement of sucrose by corn starch and increase in mono- and polyunsaturated FA by changing oil origin (flaxseed and olive oil were used instead of lard). Intervention diet was combined with an exercise training program on a treadmill (Exer-6M Open Treadmill for Mice and Rats with Shocker and Software 2-102 M/m Columbus Instruments; Columbus, OH, USA). Acclimatisation was performed already during the first week of intervention for 3 days prior 4 weeks of exercise training (5 days/week, 1 h/day). The protocol was designed to increase gradually the speed until reaching a final maximum speed of 20 m/min with a second level inclination (10°). In order to address new experimental hypothesis, a second phase of intervention was implemented for 5 extra weeks (*Int2* group). Int2 maintained the diet (100% in kcal content of the daily energy intake determined for a *Ctrl* animal) and the exercise program was adjusted to an alternate-day workout, 1 hour/day with a fixed moderate speed (14-16 m/min) and inclination (5%).

### Body composition

Mice were scanned using Magnetic Resonance Imaging in a 7.0 Teslas BioSpec 70/30 horizontal (Bruker BioSpin, Ettlingen, Germany), equipped with an actively shielded gradient system (400 mT/m, 12 cm inner diameter). Animals were placed in a supine position in a Plexiglas holder with a nose cone for administering anesthetic (1.5% isoflurane in a mixture of 30% O_2_ and 70% N_2_O). Mice were scanned from below the head to the beginning of the tail, in 1.5 mm sections. A 3D reconstruction merging all the images was performed. Fat volume shown in white contrast was calculated merging the area values for each image. A Whole-body Composition Analyser (EchoMRI™, Houston, TX, USA) was used additionally.

### Indirect calorimetry

TSE LabMaster (TSE Systems, Bad Homburg, Germany) was used as described previously (*36, 37*). Mice were acclimated for 24 h and then monitored for an additional 48h. Data was collected every 30 min for O_2_ consumption and CO_2_ production to determine energy expenditure with standard analysis software provided with the calorimeter system. Heat production was calculated by the abbreviated Weir equation ([3.94(VO_2_) + 1.11(VCO_2_)] 1.44) (*38*). Food and water intake, and locomotor activity were determined continuously for 48h. 30 min-RER values were averages for each animal in the same time point across 24 h. Median of these average for each animal was considered baseline, and the deviation to it was calculated, along with the maximal and minimum values.

### Glucose homeostasis *in vivo* functional assays

Intraperitoneal Glucose Tolerance Test (IGTT) was performed after 16 h fasting (D-Glucose at 2 g/kg of mouse BW). Blood glucose was measured before and 15, 30, 60 and 120 min after glucose administration. Insulin levels at time 0 and 15 were measured by ELISA (90080 Crystal Chem Inc., Elk Grove Village, IL, USA) according to manufacturer’s instructions. Insulin Tolerance Test (ITT) was performed after 4h fasting (insulin Humulin R 100 UI/mL Regular by Eli Lilly and Company (Indiana, IN, USA), at 0.75 UI/kg of mouse BW). Blood glucose was measured using a glucometer before and 15, 30, 60 and 120 min after insulin administration.

### *In vitrho* glucose-stimulated insulin secretion (GSIS)

Animals were anaesthetised and 2-3 mL of fresh collagenase XI solution was injected in the clamped end of the bile duct inserted in the duodenum (Collagenase from *Clostridium histolyticum*, Type XI, 2-5 FALGPA units/mg solid, >1200 CDU/mg solid; C7657 Sigma, St. Louis, MI, USA) at 1000 U/mL in Hank’s Balanced Salt Solution (HBSS without MgCl2 and CaCl2; Gibco, Thermo Fisher Scientific, Waltham, MA, USA). Pancreas tissue was then collected and incubated at 38°C for 14 min before manual agitation, and after two washes with ice-cold HBSS containing CaCl2 (HBSS, Sigma, St. Louis, MI, USA), the pellet was resuspended in 7 mL and filtered (Cell Strainer 70 um Nylon; BD Falcon, Bedford, MA, USA). The islets in the filter were washed twice with HBSS and plated in a RPMI solution by inverting the filter (RPMI, Gibco, Thermo Fisher Scientific, Waltham, MA, USA, supplemented with 2 mM L-glutamine, 1000 units/mL of both penicillin and streptomycin, Thermo Fisher Scientific, Waltham, MA, USA, and 10% FBS, Biosera, Manila, Philiphines). About 70 islets per animal were handpicked for each animal and cultured overnight in RPMI media, and then splitted in tubes of 15. To measure *insulin secretion*, islets were pre-incubated in agitation for 30 min in freshly prepared KRBH secretion solution (NaCl_2_ 140 mM, KCl 4.5 mM, MgCl_2_·6H_2_O 1mM, HEPES 20 mM, CaCl_2_ 2.5 mM, 0.1%BSA, with 2.8 mM glucose) and then incubated in KRBH solution with 2.8 mM or 16.7 mM glucose for 1h. Supernatants were collected and for *insulin content*, remaining islets were lysed in glacial acetic (5.75% (v/v) in 0.1%BSA) following an overnight freezing cycle at −80°C, warm them at 95°C for 10 min, then centrifuging them at 4°C and collecting the supernatant. Insulin levels were measured in supernatants from both *insulin secretion* and *insulin content* assays, by ELISA (Crystal Chem Inc., Elk Grove Village, IL, USA) according to manufacturer’s instructions. Insulin secretion was normalised by insulin content.

### Analytical measurements

For triglyceride measurements, lipids were extracted from tissue homogenates (100 mg of tissue in SDS 0.1%) using chloroform:methanol (2:1) to extract the organic phase pellet and resuspend it in ethanol. Triglyceride Reagent, Free Glycerol Reagent and Glycerol Standard Solution (T2449, F6428, G7793 Sigma, St. Louis, MI, USA) were used according to a modified protocol from the one provided by manufacturers in liver and skeletal muscle. Briefly, Free Glycerol Reagent was added to samples and standards in a 96-well microplate (NUNC, Denmark) incubated at 37°C for 5 min. Absorbance at 540 nm was read before and after the addition of Triglyceride Reagent (Synergy HT Multi-Mode Microplate Reader, BioTek, Winooski, VT, USA).

### Pancreas morphometry studies

Collected pancreas tissue was fixed overnight in formalin at 4°C, dehydrated and embedded in parafin blocks (Embedding Center, Leica, Buffalo Grove, IL, USA), cut in 4 μm sections separated by 150 μm (Rotary Microtome, Leica, Buffalo Grove, IL, USA) and placed in Poly-L-Lysine treated microscope slides. Paraffin was removed, and the tissue was permeabilised in PBS1X+1%Triton, blocked with 3% BSA in PBS1 and incubated overnight at 4°C with primary polyclonal antibodies for glucagon and insulin (Dako, Agilent, Santa Clara, CA, USA) (Table S8). Secondary antibodies were conjugated with Alexa Fluor®488 and Alexa Fluor®555 (Life Technologies, Carlsbad, CA, USA) (Table S8) and Hoechst was used for nuclei staining. An inverted-fluorescence microscope with camera (Leica, Buffalo Grove, IL, USA) and ImageJ were used to obtain and to measure the number and area of positive cells, respectively. The pancreas section was stained with tolonium chloride to measure the area and normalise the values.

### Gene expression

50-100 mg of tissue were homogenised in 1 mL TRI Reagent (93289 Sigma, St. Louis, MI, USA) using magnetic beads (0.5 mm diameter ZrO_2_ for BulletBlender, NextAdvance, Troy, NY, USA) in a BulletBlender following manufacturer instructions for each tissue. For adipose tissue samples, a centrifugation was performed at this point to discard the surface layer. Supernatants were collected for RNA extraction according to manufacturer’s instructions. cDNA was obtained using the High Capacity cDNA Reverse Transcription Kit (4368814 Applied Biosystems, Thermo Fisher Scientic, Waltham, MA, USA). cDNA dilution for each tissue was optimised. RT-PCR assays were performed in 384-well plates (Attendbio, Cerdanyola del Valles, Spain) using the 7900HT RT-PCR system (Thermo Fisher Scientific, Waltham, MA, USA) and the commercial reagents *Premix Ex Taq™* (BN2001 TaKaRa Bio Inc., Kusatsu, Japan) according to recommended conditions. All Taqman® probes were obtained from Applied Biosystems (Table S8). A standard curve of pooled samples was used for quantification.

### Protein content

Tissues were homogenised with magnetic beads as described above and lysed using 300 uL of lysis buffer (Na_2_HPO_4_ 10 mM, NaF 10 mM, Tris pH 7.5-8 50 mM, EDTA 5 mM, NaCl 150 mM and Triton X-100 150 mM) with the addition of a protease inhibitors cocktail (Roche), and mechanically disrupted through freeze-thaw cycles. Supernatants were collected for protein quantification and diluted at a given concentration in lysis buffer and Laemmli sample buffer 4X. Criterion XT Bis-Tris Gel (BioRad, Hercules, CA, USA) were used for electrophoresis. Transfer of the proteins to the PVDF membranes was performed using the Trans-Blot Turbo^TM^ Transfer System (BioRad, Hercules, CA, USA). Membrane was blocked for 1h at RT (5% milk in TBS-T) and incubated overnight at 4°C with primary antibodies (Table S8). Secondary antibodies were incubated for 1h at RT. Pierce^TM^ ECL Western Blotting Substrate (Thermo Fisher Scientific, Waltham, MA, USA) was used in an ImageQuant LAS 4000 (GE Healthcare, Chicago, IL, USA) for imaging of blots by chemiluminescence. ImageQuant TL Software (GE Healthcare, IL, USA) was used for the blot quantification.

### High-resolution respirometry

Mitochondrial respiration was assessed by high-resolution respirometry (HRR) in glycolytic and oxidative skeletal muscles, hypothalamus, liver and white adipose tissue. For sample preparation, each tissue was homogenised and permeabilised in fresh conditions as described in (*39*). Experiments were performed in Oxygraph-2k system (Oroboros Instruments, Innsbruck, Austria) according to an established substrate-uncoupler-inhibitor titration (SUIT) HRR protocol (https://www.mitofit.org/index.php/SUIT-008), where the nomenclature used is defined (see further details in (*35*)). Briefly, LEAK respiration was measured in the presence of the NADH (N)-linked substrates Pyruvate (5mM) and Malate (2mM). Oxidative phosphorylation capacity was then determined by the addition of ADP (5mM) at saturated concentrations, and cytochrome C (10µM) was added to assess the integrity of the mitochondrial outer membrane before the addition of Glutamate (10mM). Then, Succinate (10mM) was added to stimulate the succinate (S)-linked pathway, allowing the convergent electron flow through both pathways simultaneously. Subsequent titration of the uncoupler carbonyl cyanide p-trifluoro-methoxyphenyl hydrazine (FCCP, 0.5µM) was then performed to assess maximal non-coupled Electron Transfer (ET)-respiratory capacity mediated by NS-linked pathways. ET-respiratory capacity mediated uniquely by S-pathway was also evaluated inhibiting complex I by the addition of Rotenone (0.5µM). Finally, residual oxygen consumption (ROX) was determined by the inhibition of complex III adding Antimycin A (2,5µM) and this value was subtracted from O2 flux as a baseline for all respiratory states. The O2k-Software DatLab 7.4 was used for real-time data acquisition and analysis. Oxygen flux values were expressed relative to tissue wet weight per second (JO2, pmol mg-1 s-1).

### Adipose tissue histology

Histological examination was done using 4 μm thick sections from formalin-fixed paraffin-embedded tissue samples, which were stained with hematoxylin and eosin (H&E) at the Pathology Department of the Hospital Clinic of Barcelona. Two-four μm thickness sections from paraffin embedded samples were used for immunohistochemistry at the Tumour Bank of the HCB-IDIBAPS Biobank (IDIBAPS, Barcelona) using the Leica Microsystems’ Bond-max™ automated immunostainer together with the Bond Polymer Refine Detection System (Leica Microsystems, Spain). Briefly, tissue sections were deparaffinised, and pretreated for antigen retrieval with citrate buffer, pH 6.0 20 minutes for Caspase 3 and Mac2. For macrophage immunostaining: primary monoclonal rat anti-murine antibody to Mac2 (Cedarlane Labs, Burlington, Ontario, Canada) (Table S8) was used at 1:40,000 for one hour, combined with a secondary rabbit anti-rat Ig at 1:3,000. For apoptosis: monoclonal rabbit anti-murine Caspase3 antibody (Cell Signaling Technology, Danvers, MA, USA) (Table S8) was used at 1:500 for one hour. Finally, samples were developed with diaminobenzidine and counterstained with hematoxylin.

### Mitochondria electron microscopy

Anesthetized animals were perfused transaortically with 50 mL heparinized saline (10mg/L heparin in saline (0.9% NaCl)) followed by 100 mL of fixative solution (glutaraldehyde 2.5% + paraformaldehyde 2% in phosphate buffer 0.1 M pH=7.4). Once perfused, a small piece of adipose tissue was collected and placed on vials with fixative solution for 24 h. Samples were rinsed four times with phosphate buffer, postfixed with 1% osmium tetroxide (EMS, Hatfield, PA, USA) for 2 h and rinsed with milliQ water. Samples were then dehydrated in an acetone series (50%, 70%, 90%, 96% and 100%, for 10 min each), and infiltrated and embedded in Epon resin (EMS, Hatfield, PA, USA). Ultrathin sections of 60nm in thickness were obtained using a UC6 ultramicrotome (Leica, Buffalo Grove, IL, USA) and were stained with 2% uranyl acetate and lead citrate. Sections were observed in a Jeol 1010 (Gatan, Abitasu Yaesu, Japan) equipped with a tungsten cathode and Images were acquired at 80 kv with a CCD Megaview 1kx1k. Mitochondrial cristae density was evaluated distributing mitochondria in 4 categories: 0-not visible cristae; 1-very few cristae; 2-half of the mitochondria shows cristae, 3-equaly distributed cristae.

### Whole mount adipose tissue immunostaining and vessel area quantification

Approximately 3 mm^3^ cubes of adipose tissue were cut from each sample and permeabilised for 1h with phosphate buffer saline (PBS) and 1% Triton-x 100. Afterwards, tissues were blocked with blocking buffer (PBS + 0,3% Triton-x 100 + 5% goat serum) for 2 h at room temperature. Primary antibody (Isolectin GS-B4 568, Invitrogen, Thermo Fisher Scientic, Waltham, MA, USA) (Table S8) diluted (1:2000) in blocking buffer was incubated overnight at room temperature, with over day washings with PBS + 0,3% Triton-x 100. Afterwards, samples were clarified using 90% glycerol at 4°C overnight. Samples were stored in ProLong™ Gold Antifade Reagent (P36930 Invitrogen, Thermo Fisher Scientic, Waltham, MA, USA) with DAPI. Images were taken with a Leica SP5 confocal microscope using 20X and 40X objectives (Leica, Buffalo Grove, IL, USA). All images are maximal z-stack projections. Images were processed using Volocity, Fiji and Adobe Photoshop CS5. Vessel density was quantified by measuring isolectin-B4 positive area, within a fixed square template, using the ImageJ software.

### Phospholipid extraction

White adipose tissue (WAT) was homogenised in PBS of which 5% was removed to determine protein content via Bradford assay. Next, lipids were extracted twice from homogenate with 2:1 chloroform:methanol (Folch method) containing 0.5 μM CL(14:0)_4_ as internal standard and dried under N_2_ flow. Due to the high content of neutral lipids in WAT, phospholipids were separated by solid-phase extraction using Sep-Pak C18-columns (Waters Associates, Milford, MA, USA). Briefly, the lipid extract was re-suspended in 1 mL chloroform, applied on the column and washed thrice with 1 mL chloroform to elute neutral lipids. Phospholipids were eluted by quadruple addition of 2:1 chloroform:methanol and quadruple addition of methanol, collected and dried under N_2_ flow.

### LC-MS/MS analysis of CL MLCL

CL analysis was performed as described in (*40*). Phospholipid extracts were dissolved in HPLC starting condition and subjected to HPLC-MS/MS analysis. Separation was achieved by reversed-phase HPLC with an Agilent Poroshell 120 EC-C8 2.7 μm 2.1×100mm column (Agilent Technologies, Santa Clara, CA, USA) on a Dionex Ultimate 3000 HPLC (Thermo Fisher Scientific, Waltham, MA, USA, 50°C column oven, 0.4 mL/min flow) with running solvent A (60/40 Acetonitrile/H_2_O, 10 mM ammonium formate, 0.2% formic acid) and running solvent B (90/10 Isopropanol/Acetonitrile, 10 mM ammonium formate, 0.2% formic acid). Analytes were measured using a LTQ Velos MS (Thermo Fisher Scientific, Waltham, MA, USA) operated in negative ESI mode (3.8kV, 275°C capillary temperature, 460-1650 m/z) and data-dependent MS2 acquisition. Thermo raw data was converted to open-source MZML format and Peaks were integrated in MZmine2 (*41*). Identification was based on a combination of accurate mass, (relative) retention times, and fragmentation spectra, compared to a library of standards. Data was corrected for internal standard, normalised on protein content, quantified using an external dilution series (CL(14:0)_4_, CL(18:1)_4_) and further analysed by an in-house pipeline in R. Significance was calculated using a one-way ANOVA with post hoc Tukey correction.

### Quantification of mtDNA copy number

Mitochondrial DNA (mtDNA) content was determined by qPCR of total DNA extracted. NucleoSpin Tissue DNA purification kit (Macherey-Nagel, Dueren, Germany) was used to extract total DNA from adipose tissue according to manufacturer’s protocol, and NanoDrop 1000 Spectrophotometer (Thermo Fisher Scientific, Waltham, MA, USA) to quantify DNA concentration. GoTaq® qPCR MasterMix (Promega, Madison, WI, USA) and 3ng of DNA sample were combined to amplify both mtDNA and nuclear DNA by 7900HT RT-PCR system (Thermo Fisher Scientific, Waltham, MA, USA) using the following primers; cytochrome C subunit 2 (*mt-Co2* Fw: CTACAAGACGCCACAT – *mt-Co2* Rv: GAGAGGGGAGAGCAAT), 12S rRNA (*Mt-Rnr1*Fw: CTTCAGCAAACCCTAAAAAGG – *Mt-Rnr1*Rv: GCGTACTTCATTGCTCAATTC) and succinate dehydrogenase subunit A (*Sdha* Fw: TACTACAGCCCCAAGTCT - *Sdha* Rv: TGGACCCATCTTCTATGC). mtDNA content is referred to nuclear DNA (nDNA) as the copy number ratio of mtDNA to nDNA.

### Metabolomics sample preparation

Pulverised liver tissue (50-100 mg) was mixed with ice-cold acetonitrile:water (1:1) and ultrasonicated, and the combined supernatant aqueous phases of 3 repetitions were frozen and lyophilised. The resultant pellet was dried and extracted with chloroform:methanol (2:1) by ultrasonication. The supernatant organic phase was collected and dried under N_2_ stream. Pulverised white adipose (30-70 mg), gastrocnemius (20-40 mg) and hypothalamus (10-25 mg) tissue was mixed with methanol and ultrasonicated before chloroform was added in two steps to final 1:1. Water was added 1:2:2 (water:methanol:chloroform) and the aqueous upper phase containing water and methanol and the organic lower phase were collected separately. An aqueous extract was dissolved in D2O (containing 0.67 mM trisilylpropionic, TSP) and the organic extract was reconstituted in CDCl_3_/CD_3_OD (2:1) solution (containing 1.18 mM tetramethylsilane, TMS). The supernatant of both phases was transferred into 5mm NMR tubes.

### Nuclear Magnetic Ressonance (NMR) metabolomics analysis

^1^H-NMR spectra were recorded at 310K on a Bruker Avance III 600 spectrometer (Bruker, Billerica, USA) operating at a proton frequency of 600.20MHz using a 5mm CPTCI triple resonance (^1^H, ^13^C, ^31^P). One-dimensional ^1^H pulse experiments of aqueous samples were performed. In order to suppress the residual water peak the experiments were recorded using the *NOESY-presat* sequence, with the mixing time set at 100ms. A 75Hz-power irradiation was applied during recycling delay and mixing time to presaturate the solvent. A total of 256 transients were collected into 64 k data points for each ^1^H spectrum, being the spectral width set as 12 kHz (20 ppm). The exponential line broadening applied before Fourier transformation was of 0.3Hz. In order to remove the residual water moisture of deuterated methanol, lipidic samples were measured using a simple presaturation sequence. Thus, a 50Hz-power irradiation was used during recycling delay and mixing time to presaturate the solvent. Again, 256 transients were collected into 64 k data points for each ^1^H spectrum, being the spectral width set as 12kHz (20 ppm). The exponential line broadening applied before Fourier transformation was also here of 0.3Hz. Of note, the frequency domain spectra were manually phased and baseline-corrected using TopSpin software (version 2.1, Bruker, Billerica, USA).

### NMR data analysis

All the acquired ^1^H NMR spectra were phased, baseline-corrected, and referenced to the chemical shift of TSP or TMS signal at 0 ppm. For metabolic identification, references of pure compounds from the metabolic profiling AMIX spectra database (Bruker, Billerica, USA), human metabolome database (HMDB), and Chenomx databases were used (Chenomx Inc. Edmonton, Canada). After baseline correction, specific ^1^H NMR regions identified in the spectra were integrated for each extraction method entering the study using the AMIX 3.8 software package. Then, each integration region was normalised using the tissue weight used from each sample. Data (pre-) processing, data analysis, and statistical calculations were performed in RStudio (R version 3.0.2).

### RNA-Seq data processing and differential expression analysis

RNA samples from 20 WAT samples (Ctrl n=8, HFD n=4, Int n=4 and Int2 n=4) as well as 12 liver tissue samples (Ctrl n=4, HFD n=4 and Int n=4) were sequenced by standard Illumina protocol to create raw sequence files (.fastq files), which underwent quality control analysis using FastQC (http://www.bioinformatics.babraham.ac.uk/projects/fastqc/). We aligned the quality checked reads to the Mouse Genome, version July 2007 (NCBI37/mm9) using TopHat version 2.1.1 allowing for unique alignments to the genome and up to two mismatches. The resulting alignments were summarised by Ensembl gene identifiers to evaluate number of uniquely aligned reads per transcript and per sample (raw read counts).

RNA-seq data were analysed using the limma package version 1.8.2 available through the Bioconductor open source. The raw read counts were used as input to form a DGEList object combining the counts and associated annotation. Scale normalisation was applied and the calculation of normalised signal was performed by voom function of the limma package (*42*) available through the Bioconductor open source software. This differential expression analysis was performed pair-wise between different experimental conditions, but per each tissue type separately (FDR<0.05). First, we identified genes either up- or downregulated in HFD conditions as compared to controls. Then, we explored the expression modulation of these two sets of genes after first intervention (Int). We could determine three distinct patterns: 1) genes fully recovered (with differential expression between HFD and Int but no significant difference in expression between controls and Int); 2) genes partially recovered (not reaching the same expression level as in control conditions but showing the recovering tendency between HFD and Int); 3) genes not recovered (that continue to be differentially expressed between Int and controls, with no significant difference in expression between HFD and Int). In the case of WAT, for the third, not recovered category, we performed similar expression patterns analysis using the data of second intervention (Int2), defining the genes that recovered or not after this additional experimental step.

### Gene Ontology analysis

For different gene sets, GO enrichments were performed using the GOstats R package (*43*), separately for biological processes and cell compartments related GO terms. Significant GO terms (p<0.0001) were further summarised together by ReVigo software (*44*) in order to represent them as semantic-similarity scatter plot of the most significant GO terms, with a modification of the script to represent the GO enrichment values as dot size. To represent the cascade of GO terms affected in different steps of the experiment (HFD, Int and Int2, Figure5), we combined a biological and semantic approach, merging those GO terms that contained the same genes and that were related to the same biological process or cell compartment. In this analysis, general, ancestor GO terms (the highest level in GO terms tree; typically containing over 500 genes) as well as very specific GO terms (typically containing less than 5 genes) were excluded, in order to obtain a clearer view of the GO terms dynamics throughout three different experimental conditions.

### Database and pattern data analysis

Gene expression (transcriptomics), metabolites content (metabolomics) and mitochondrial respiration states (high-resolution respirometry) patterns were analysed and are presented in Figure 2. A Content Management System (CMS) dynamic database with a front-end user interface or a content management application (CMA) (idibaps.seeslab.net) was created along with a Content Delivery Application (CDA) that compiles all the information and updates the website. An entry for each mouse was generated including both *phenotypical information* (e.g., animal identification, group (Ctrl, HFD or Int), body weight, glucose levels (IGTT, ITT), etc) and *specific-tissue data* (e.g., mitochondrial respiratory values, gene expression data, protein content data, targeted metabolomics data, etc). Python scripts allowed correlation and comparison analysis of all the attributes entered in the database, which were structured in a pandas. DataFrame (http://pandas.pydata.org/) generated from the database. NumPy and SciPy were used for working with multi-dimensional arrays and matrices as well as for calculating high-level mathematical functions. Matplotlib was used for graphical representations (i.e., stacked bar plot). Pattern data analysis: to visually show the proportion of parameters that were reversed after the intervention we defined a 3-point vector for each parameter (e.g., gene expression, metabolite, mitochondrial respiration state) in which the 3 points correlated to the mean of this given parameter for 1) Ctrl group 1) HFD-fed group and 3) Int group (i.e., *parameter=[Ctrl mean, HFD mean, Int mean]*). Mann-Whitney comparisons were used to compare the groups. Once the significant differences were calculated, different patterns were defined: green for a reversion pattern after the intervention; red for no reversion in the intervention group, so the mean value in the HFD group, which is significantly different from the Ctrl group, is not different in comparison to the Int group; and grey for a difference in the Int group in comparison to the other two groups, between the ones there is no difference. Once these patterns were defined for gene expression, metabolites and mitochondrial respiration states in skeletal muscle (gastrocnemius, soleus and tibialis anterior), hypothalamus, liver and epididymal adipose tissue, stacked bar plots were used to visualise the relative percentage of all the genes/metabolites/respiratory states falling in the different patterns.

### Statistical analysis

Results are expressed as Mean±SEM. The statistical significance among the three experimental groups was assessed using one-way ANOVA, and differences between means were subsequently tested by Tukey post-hoc test. Data curation involved the removal of values that were ±2 standard deviation (SD) in punctual situations. A p-value <0.05 was considered significant in all cases, meaning a confidence interval of 95% and setting significance level at α=0.05. Tendencies with p-value between 0.05 and 0.07 are also indicated. All statistical analyses were performed using GraphPad 6 (GraphPad Software, Inc. La Jolla, CA, USA).

### Lead Contact

Further information and requests for resources should be directed to and will be fulfilled by the Lead Contact, Pablo M. Garcia-Roves.

## Supplementary information

**Supplementary Figure 1.**
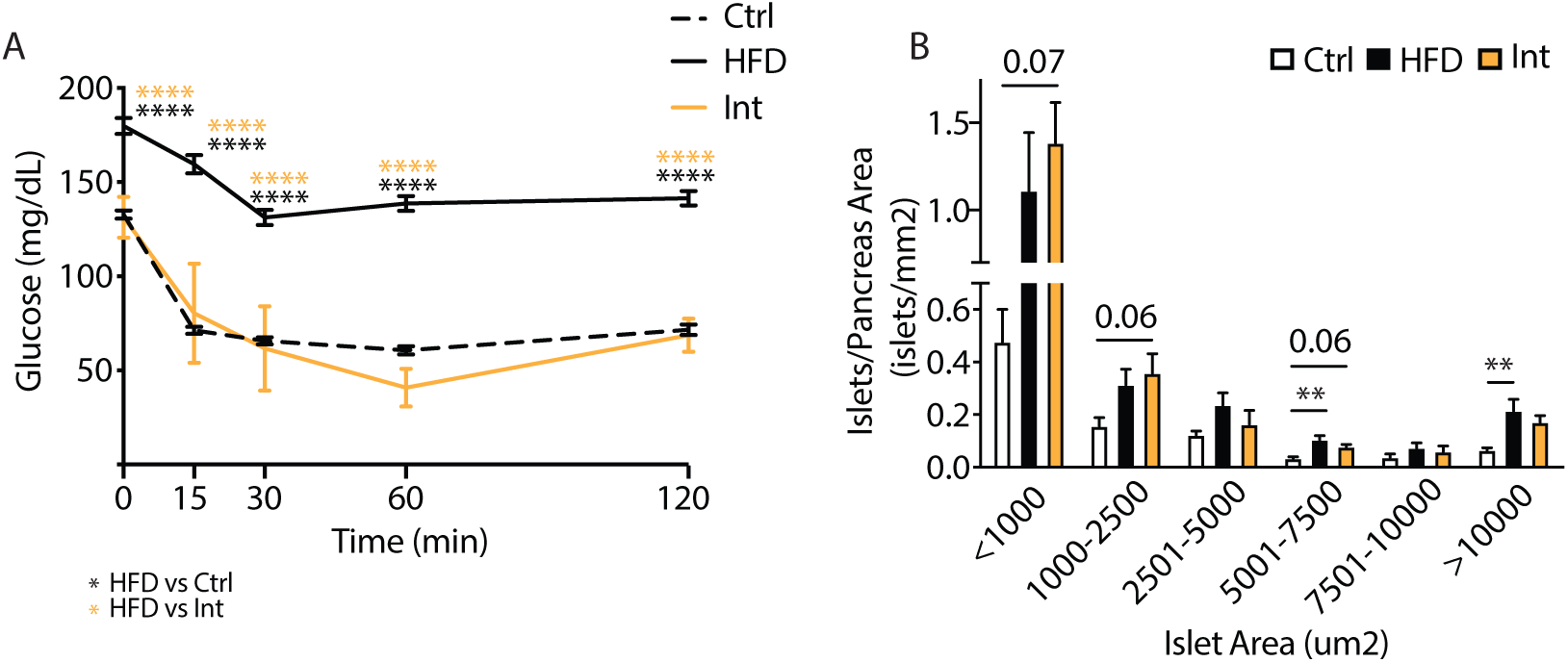
A: Insulin tolerance test in fasted mice (Ctrl n=9; HFD n=6; Int n=6). B: Beta-cell size distribution per pancreas area (Ctrl n=6; HFD n=4; Int n=4). Data represented as mean ± SEM, ANOVA One-way and Post-hoc Tukey, *p<0.05, **p<0.01, ***p<0.001, and ****p<0.0001.

**Supplementary Figure 2.**
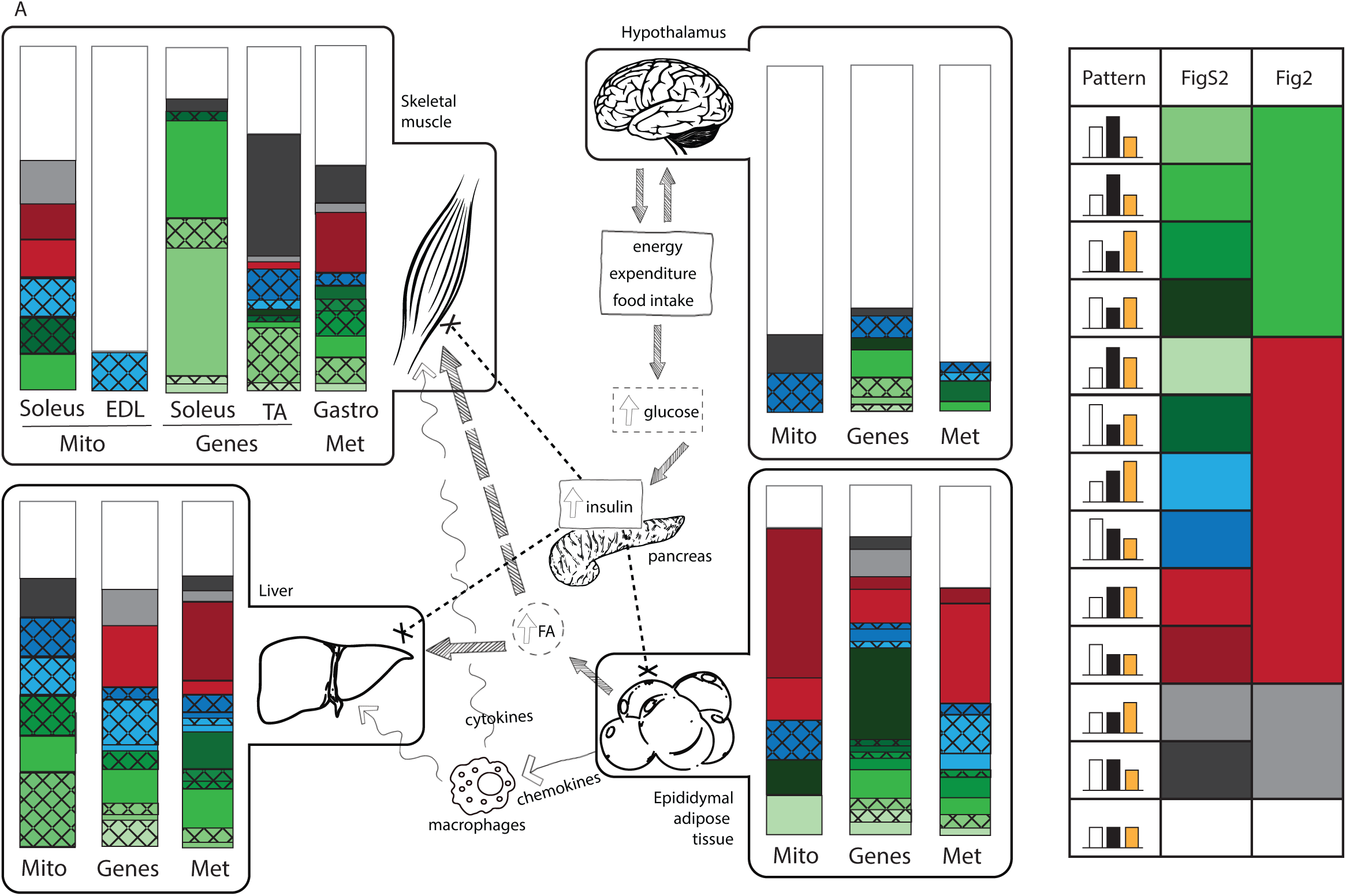
A: Detailed representation of Figure 2A uncovering sub-patterns within each group. Different colors indicate the different sub-patterns (in FigureS2A) that correspond to patterns (in Figure 2A). Patterns are grouped in 4 different colors for Figure 2A for simplification. A crossing pattern in Figure S2A indicates no transitivity property, in 3 scenarios: Ctrl vs HFD with no statistical difference, Ctrl vs Int with no statistical difference, but HFD vs Int with a statistical difference (Ctrl≈HFD, Ctrl≈Int, HFD≠Int); when Ctrl≈HFD, Ctrl≠Int, HFD≈Int; and when Ctrl≠HFD, Ctrl≈Int, HFD≈Int.

**Supplementary Figure 3.**
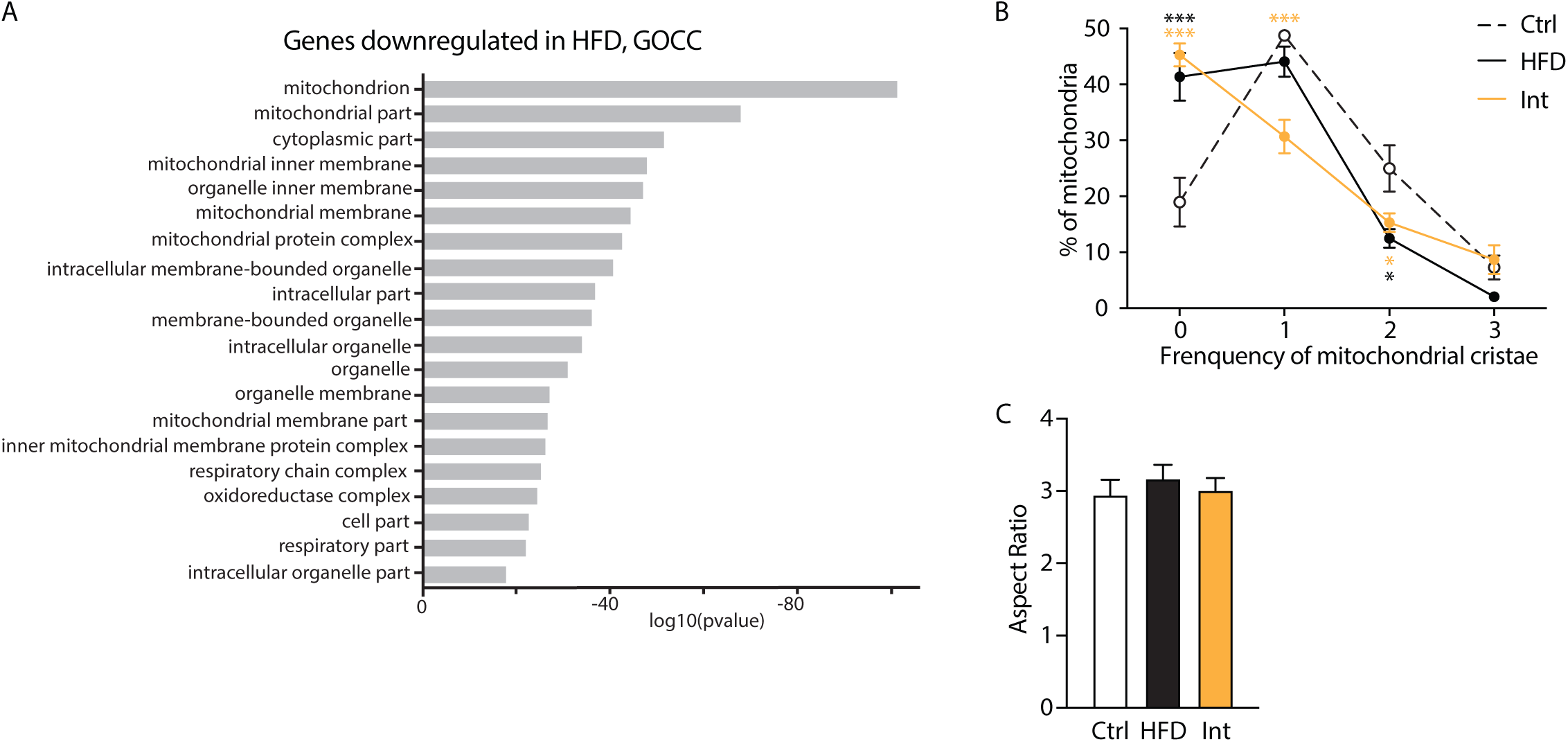
A: Gene ontology cellular components (GOCC) enrichment analysis for the 1369 transcripts downregulated in HFD and Int. B: Percentage of mitochondria within the different mitochondrial cristae frequency categories (Ctrl n=3, HFD n=4, Int n=4). C: Mitochondrial aspect ratio (Ctrl n=3, HFD n=4, Int n=4).

**Supplementary Figure 4.**
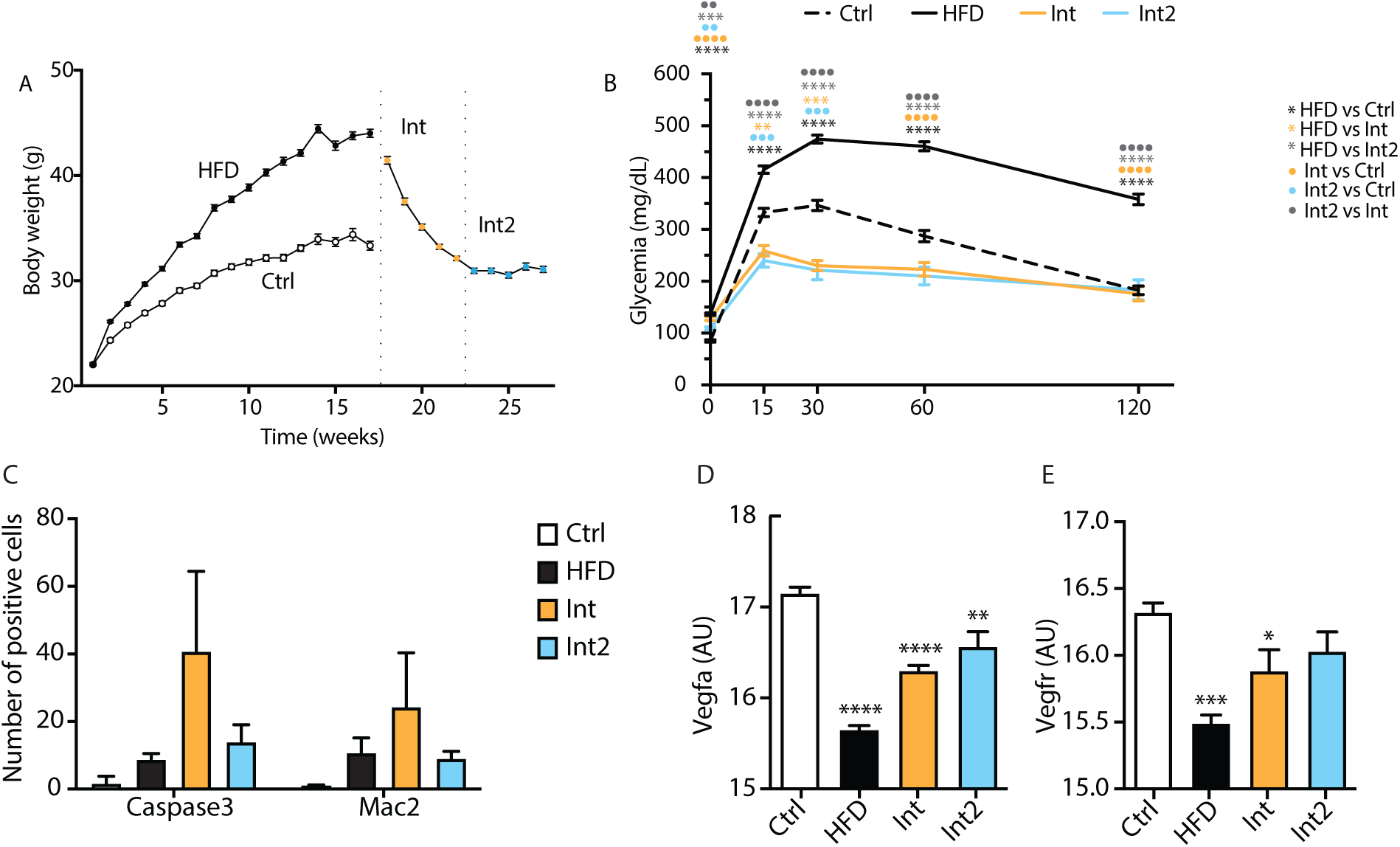
A: Body weight evolution in the 4 experimental groups. B: Intraperitoneal glucose tolerance test (IGTT) (Ctrl n=82, HFD n=168, Int n=32, Int2 n=19). C: Quantification for Caspase3- and Mac2-positive cells staining in eWAT for each experimental group (Figure 4D) (Ctrl n=3-4, HFD n=4, Int n=3-4, Int2 n=4-5). D and E: Vegfa (c) and Vegfr (d) normalized eWAT gene expression (Ctrl n=8, HFD n=4, Int n=4, Int2 n=4). Ctrl: control; HFD: high fat diet-fed; Int: intervention; Int2: second intervention. Data represented as mean ± SEM, ANOVA One-way and Post-hoc Tukey, significant differences with respect to Ctrl *p<0.05, **p<0.01, ***p<0.001, and ****p<0.0001.

### Supplementary Tables 1-7

http://resources.idibaps.org/paper/obesity-triggers-a-progressive-mitochondrial-deterioration-in-white-adipose-tissue-even-after-a-lifestyle-mediated-phenotypic-reversion

**Supplementary Table 1**. Gene expression: Gene expression by RT-PCR in the different tissues and experimental groups (corresponding to Figure 2).

https://drive.google.com/open?id=1wArleHoqkSYAnjBa2pRruNz9ytSfG15T

**Supplementary Table 2**. Gene expression: Gene expression by RNA-seq in the different tissues and experimental groups (corresponding to Figures 2-4).

https://drive.google.com/open?id=1hyla2k2TEFf3zEohIzQRfCnFaCMxGamr

**Supplementary Table 3.** Metabolomics: Metabolic data for the different tissues and experimental groups (corresponding to Figure 2).

https://drive.google.com/open?id=17hULD46zDsrPFnDdH1sYFfTafW A7MQ

**Supplementary Table 4.** Mitochondria function: High-resolution respirometry of the different tissues and experimental groups (corresponding to Figures 2-4).

https://drive.google.com/open?id=1D39FCgM3-npu-CZq_ZLTwNn-h8pN05OZ

**Supplementary Table 5.** Cardiolipidomics: Cardiolipin anaylsis by liquid chromatography with mass spectrometry for the different experimental groups (Corresponding to Figure 5).

https://drive.google.com/open?id=1rcE8PfkhevjLB5_wRJEeFyznjh4q_1WJ

**Supplementary Table 6.** Gene ontologies of genes irreversibly decreased with HFD-feeding: Gene ontology biological processes enrichment analysis in the clusters of genes irreversibly decreased with HFD-feeding; from the first intervention and form the second intervention (corresponding to Figure 5).

https://drive.google.com/open?id=1zgEHbjyZvI_ux4VId7zTeAnKXzH_9JHI

**Supplementary Table 7**. Gene ontologies of genes irreversibly decreased with HFD-feeding: Gene ontology cellular compartments enrichment analysis in the clusters of genes irreversibly decreased with HFD-feeding; from the first intervention and form the second intervention (corresponding to Figure 5).

https://drive.google.com/open?id=1z9VmKXBxGEgiwB3KbOtQ4SyKc_ZF5LMY

**Supplementary Table 8.** Antibodies, TaqMan probes and mitochondrial substrates, uncoupler and inhibitors (Mitochondrial High-Resolution Respirometry).

|  | Source | Reference |
| --- | --- | --- |
| **Antibodies** |  |  |
| Polyclonal Rabbit Anti-Human Glucagon antibody, 1:1000 | Dako / Agilent | A0565 |
| Polyclonal Guinea Pig Anti- Insulin antibody, 1:1000 | Dako / Agilent | A0564 |
| Donkey Anti-Rabbit IgG (H+L) Antibody, Alexa Fluor 488 Conjugated, 1:500 | Life Technologies | A21206 |
| Goat Anti-Guinea Pig IgG Antibody, Alexa Fluor 555 Conjugated, 1:500 | Life Technologies | A21435 |
| Mouse Anti-OPA1 Monoclonal Antibody, Unconjugated, Clone 18, 1:1000 | BD Biosciences | 612606 |
| Rat Anti-Mouse MAC-2 and Gal-3 (Galectin-3) Monoclonal antibody, Unconjugated, Clone m3/38, 1:40000 | CEDARLANE | CL8942AP |
| Cleaved Caspase-3 (Asp175) (5A1E) Rabbit mAb antibody, 1:500 | Cell Signaling Technology | 9664 |
| Isolectin GS-IB<sub>4</sub> From Griffonia simplicifolia, Alexa Fluor™ 568 Conjugate, 1:2000 | Invitrogen | I21412 |
| DAPI |  |  |
| High-resolution respirometry: substrates, inhibitors and uncoupler |  |  |
| L-Glutamic acid, sodium salt, C<sub>5</sub>H<sub>8</sub>NO<sub>4</sub>Na | Sigma | G1626 |
| L-Malic acid, C<sub>4</sub>H<sub>6</sub>O<sub>5</sub> | Sigma | M1000 |
| Pyruvic acid sodium salt, C<sub>3</sub>H<sub>3</sub>O<sub>3</sub>Na | Sigma | P2256 |
| Succinate disodium salt, hexahydrate, C<sub>4</sub>H<sub>4</sub>O<sub>4</sub>Na<sub>2</sub> x 6 H<sub>2</sub>O | Sigma | S2378 |
| Cytochrome c | Sigma | C7752 |
| Adenosine 5'diphosphate, C<sub>10</sub>H<sub>15</sub>N<sub>5</sub>O<sub>10</sub>P<sub>2</sub>K, potassium salt, contains 1 mol/mol H<sub>2</sub>O, ADP | Sigma | A5285 |
| Carbonyl cyanide p- (trifluoro-methoxy) phenyl-hydrazine C<sub>10</sub>H<sub>5</sub>F<sub>3</sub>N<sub>4</sub>O, FCCP | Sigma | C2920 |
| Antimycin A | Sigma | A8674 |
| Rotenone, C<sub>23</sub>H<sub>22</sub>O<sub>6</sub> | Sigma | R8875 |
| Taqman® probes |  |  |
| acetyl-Coenzyme A carboxylase alpha | Acaca | Mm01304257_m1 |
| adhesion G protein-coupled receptor E1 | Adgre1 | Mm00802529_m1 |
| agouti related neuropeptide | Agrp | Mm00475829_g1 |
| activating transcription factor 4 | Atf4 | Mm00515324-m1 |
| activating transcription factor 6 | Atf6 | Mm01295317-m1 |
| ATP synthase, H<sup>+</sup> transporting, mitochondrial F1 complex, alpha subunit 1 | Atp5a1 | Mm00431960-m1 |
| CART prepropeptide | Cartpt | Mm04210469_m1 |
| catalase | Cat | Mm00437992-m1 |
| chemokine (C-C motif) ligand 2 | Ccl2 | Mm00441242_m1 |
| CD209e antigen | Cd209e | Mm00459980_m1 |
| CD68 antigen | Cd68 | Mm00839636_g1 |
| cell death-inducing DNA fragmentation factor, alpha subunit-like effector A | Cidea | Mm00432554_m1 |
| caseinolytic mitochondrial matrix peptidase chaperone subunit | Clpx | Mm00488586_m1 |
| cytochrome c oxidase subunit IV isoform 1 | Cox4i1 | Mm01250094-m1 |
| carnitine palmitoyltransferase 1a, liver | Cpt1a | Mm00550438-m1 |
| corticotropin releasing hormone | Crh | Mm01293920_s1 |
| citrate synthase | Cs | Mm00466043-m1 |
| DNA-damage inducible transcript 3 | Ddit3 | Mm00492097_m1 |
| diacylglycerol O-acyltransferase 2 | Dgat2 | Mm00499536_m1 |
| dynamamin 1-like | Dnm1l | Mm01342903_m1 |
| estrogen related receptor, alpha | Esrra | Mm00433143-m1 |
| fatty acid synthase | Fasn | Mm00662319-m1 |
| fibroblast growth factor 1 | Fgf1 | Mm00438906_m1 |
| fibroblast growth factor 21 | Fgf21 | Mm00840165_g1 |
| fission 1 (mitochondrial outer membrane) | Fis1 | Mm00481580_m1 |
| forkhead box O1 | Foxo1 | Mm00490672_m1 |
| GA repeat binding protein, alpha | Gabpa | Mm00484598-m1 |
| glyceraldehyde-3-phosphate dehydrogenase | Gapdh | Mm99999915_g1 |
| glutathione peroxidase 1 | Gpx1 | Mm00656767-g1 |
| glycogen synthase 1, muscle | Gys1 | Mm01962575_s1 |
| glycogen synthase 2 | Gys2 | Mm00523953_m1 |
| hepatic nuclear factor 4, alpha | Hnf4a | Mm01247716_m1 |
| heat shock protein 5 | Hspa5 | Mm00517691-m1 |
| heat shock protein 9 | Hspa9 | Mm00477716_g1 |
| heat shock protein 1 (chaperonin) | Hspd1 | Mm00849835_g1 |
| interleukin 10 | IL10 | Mm00439614_m1 |
| interleukin 1 beta | IL1b | Mm00434228_m1 |
| interleukin 6 | IL6 | Mm00446190_m1 |
| integrin alpha X | Itgax | Mm00498698_m1 |
| leptin receptor | Lepr | Mm00440181_m1 |
| lon peptidase 1, mitochondrial | Lonp1 | Mm01236887_m1 |
| lipoprotein lipase | Lpl | Mm00434764_m1 |
| melanocortin 3 receptor | Mc3r | Mm00434876_s1 |
| melanocortin 4 receptor | Mc4r | Mm00457483_s1 |
| mitofusin 1 | Mfn1 | Mm00612599_m1 |
| mitofusin 2 | Mfn2 | Mm00500120_m1 |
| mitochondrially encoded cytochrome c oxidase I | mt-Co1 | Mm04225243_g1 |
| mitochondrially encoded cytochrome b | mt-Cytb | Mm04225271_g1 |
| mitochondrially encoded NADH dehydrogenase 3 | mt-Nd3 | Mm04225292_g1 |
| NADH dehydrogenase (ubiquinone) 1 alpha subcomplex, 9 | Ndufa9 | Mm00481216-m1 |
| nicotinamide N-methyltransferase | Nnmt | Mm00447994_m1 |
| neuropeptide Y | Npy | Mm00445771_m1 |
| nuclear respiratory factor 1 | Nrf1 | Mm00447996_m1 |
| optic atrophy 1 | Opa1 | Mm00453879_m1 |
| phosphoenolpyruvate carboxykinase 1, cytosolic | Pck1 | Mm00440636_m1 |
| phosphofructokinase, liver, B-type | Pfkl | Mm00435587_m1 |
| phosphofructokinase, muscle | Pfkm | Mm01309576_m1 |
| pro-opiomelanocortin-alpha | Pomc | Mm00435874_m1 |
| peroxisome proliferator activated receptor alpha | Ppara | Mm00440939_m1 |
| peroxisome proliferator activator receptor delta | Ppard | Mm00803184-m1 |
| peroxisome proliferator activated receptor gamma | Pparg | Mm00440945_m1 |
| peroxisome proliferative activated receptor gamma coactivator 1 alpha | Ppargc1a | Mm01208835-m1 |
| peroxisome proliferative activated recetor gamma coactivator 1 beta | Ppargc1b | Mm01258518_m1 |
| peptidylprolyl isomerase A | Ppia | Mm02342430_g1 |
| stearoyl-Coenzyme A desaturase 1 | Scd1 | Mm01197142-m1 |
| succinate dehydrogenase complex, subunit A, flavoprotein (Fp) | Sdha | Mm01352366-m1 |
| solute carrier family 2 (facilitated glucose transporter), member 2 | Slc2a2 | Mm00446224_m1 |
| solute carrier family 2 (facilitated glucose transporter), member 4 | Slc2a4 | Mm00436615_m1 |
| suppressor of cytokine signalling 3 | Socs3 | Mm00545913_s1 |
| superoxide dismutase 2, mitochondrial | Sod2 | Mm01313000-m1 |
| sterol regulatory element binding transcription factor 1 | Srebf1 | Mm00550338_m1 |
| transcription factor A, mitochondrial | Tfam | Mm00447485-m1 |
| tumor necrosis factor | Tnf | Mm00443258_m1 |
| ubiquitin-like 5 | Ubl5 | Mm01151181_m1 |
| uncoupling protein 1 (mitochondrial, proton carrier) | Ucp1 | Mm00494069_m1 |
| uncoupling protein 2 (mitochondrial, proton carrier) | Ucp2 | Mm00627599_m1 |
| uncoupling protein 3 (mitochondrial, proton carrier) | Ucp3 | Mm00494077_m1 |
| ubiquinol cytochrome c reductase core protein 2 | Uqcrc2 | Mm00445961-m1 |
| X-box binding protein 1 spliced | Xbp1sp | Mm03464496_m1 |
| X-box binding protein 1 total | Xbp1t | Mm00457357_m1 |

## References

1. WHO, https://www.who.int/en/news-room/fact-sheets/detail/obesity-and-overweight.

2. M. Bluher, Obesity: global epidemiology and pathogenesis. Nat Rev Endocrinol 15, 288–298 (2019).

3. M. A. van Baak, E. C. M. Mariman, Mechanisms of weight regain after weight loss - the role of adipose tissue. Nat Rev Endocrinol 15, 274–287 (2019).

4. S. B. Heymsfield, T. A. Wadden, Mechanisms, Pathophysiology, and Management of Obesity. N Engl J Med 376, 1492 (2017).

5. F. Bäckhed et al., The next decade of metabolism. Nature Metabolism 1, 2–4 (2019).

6. DIAMAP, https://www.diamap.eu/roadmap/domain/detail/3.

7. V. A. Fonseca, Defining and characterizing the progression of type 2 diabetes. Diabetes Care 32 Suppl 2, S151–156 (2009).

8. J. E. Galgani, C. Moro, E. Ravussin, Metabolic flexibility and insulin resistance. Am J Physiol Endocrinol Metab 295, E1009–1017 (2008).

9. L. Wilson-Fritch et al., Mitochondrial remodeling in adipose tissue associated with obesity and treatment with rosiglitazone. J Clin Invest 114, 1281–1289 (2004).

10. J. X. Rong et al., Adipose mitochondrial biogenesis is suppressed in db/db and high-fat diet-fed mice and improved by rosiglitazone. Diabetes 56, 1751–1760 (2007).

11. A. De Pauw, S. Tejerina, M. Raes, J. Keijer, T. Arnould, Mitochondrial (dys)function in adipocyte (de)differentiation and systemic metabolic alterations. Am J Pathol 175, 927–939 (2009).

12. S. Boudina, T. E. Graham, Mitochondrial function/dysfunction in white adipose tissue. Exp Physiol 99, 1168–1178 (2014).

13. J. H. Lee et al., The Role of Adipose Tissue Mitochondria: Regulation of Mitochondrial Function for the Treatment of Metabolic Diseases. Int J Mol Sci 20, (2019).

14. S. Carobbio, V. Pellegrinelli, A. Vidal-Puig, Adipose Tissue Function and Expandability as Determinants of Lipotoxicity and the Metabolic Syndrome. Adv Exp Med Biol 960, 161–196 (2017).

15. K. Sun et al., Dichotomous effects of VEGF-A on adipose tissue dysfunction. Proc Natl Acad Sci U S A 109, 5874–5879 (2012).

16. I. Wernstedt Asterholm et al., Adipocyte inflammation is essential for healthy adipose tissue expansion and remodeling. Cell Metab 20, 103–118 (2014).

17. C. Crewe, Y. A. An, P. E. Scherer, The ominous triad of adipose tissue dysfunction: inflammation, fibrosis, and impaired angiogenesis. J Clin Invest 127, 74–82 (2017).

18. F. P. Hoevenaars et al., Adipose tissue metabolism and inflammation are differently affected by weight loss in obese mice due to either a high-fat diet restriction or change to a low-fat diet. Genes Nutr 9, 391 (2014).

19. D. Y. Jung et al., Short-term weight loss attenuates local tissue inflammation and improves insulin sensitivity without affecting adipose inflammation in obese mice. Am J Physiol Endocrinol Metab 304, E964–976 (2013).

20. A. Kosteli et al., Weight loss and lipolysis promote a dynamic immune response in murine adipose tissue. J Clin Invest 120, 3466–3479 (2010).

21. B. Liu et al., Intermittent Fasting Improves Glucose Tolerance and Promotes Adipose Tissue Remodeling in Male Mice Fed a High-Fat Diet. Endocrinology 160, 169–180 (2019).

22. J. Schmitz et al., Obesogenic memory can confer long-term increases in adipose tissue but not liver inflammation and insulin resistance after weight loss. Mol Metab 5, 328–339 (2016).

23. B. F. Zamarron et al., Macrophage Proliferation Sustains Adipose Tissue Inflammation in Formerly Obese Mice. Diabetes 66, 392–406 (2017).

24. S. Michel et al., Crosstalk between mitochondrial (dys)function and mitochondrial abundance. J Cell Physiol 227, 2297–2310 (2012).

25. F. Boos, J. Labbadia, J. M. Herrmann, How the Mitoprotein-Induced Stress Response Safeguards the Cytosol: A Unified View. Trends Cell Biol 30, 241–254 (2020).

26. Y. A. An et al., Dysregulation of Amyloid Precursor Protein Impairs Adipose Tissue Mitochondrial Function and Promotes Obesity. Nat Metab 1, 1243–1257 (2019).

27. A. J. Chicco, G. C. Sparagna, Role of cardiolipin alterations in mitochondrial dysfunction and disease. Am J Physiol Cell Physiol 292, C33–44 (2007).

28. G. Oemer et al., Phospholipid Acyl Chain Diversity Controls the Tissue-Specific Assembly of Mitochondrial Cardiolipins. Cell Rep 30, 4281–4291 e4284 (2020).

29. J. Li et al., Cardiolipin remodeling by ALCAT1 links oxidative stress and mitochondrial dysfunction to obesity. Cell Metab 12, 154–165 (2010).

30. X. Liu et al., Ablation of ALCAT1 mitigates hypertrophic cardiomyopathy through effects on oxidative stress and mitophagy. Mol Cell Biol 32, 4493–4504 (2012).

31. C. Song et al., Cardiolipin remodeling by ALCAT1 links mitochondrial dysfunction to Parkinson’s diseases. Aging Cell 18, e12941 (2019).

32. O. Hahn et al., A nutritional memory effect counteracts benefits of dietary restriction in old mice. Nat Metab 1, 1059–1073 (2019).

33. S. Gumeni, I. P. Trougakos, Cross Talk of Proteostasis and Mitostasis in Cellular Homeodynamics, Ageing, and Disease. Oxid Med Cell Longev 2016, 4587691 (2016).

34. C. Lopez-Otin, M. A. Blasco, L. Partridge, M. Serrano, G. Kroemer, The hallmarks of aging. Cell 153, 1194–1217 (2013).

35. E. Gnaiger, Mitochondrial respiratory states and rates. MitoFit Preprint Arch 10.26124/mitofit:190001.v6, (2019).

36. P. T. Pfluger et al., Simultaneous deletion of ghrelin and its receptor increases motor activity and energy expenditure. Am J Physiol Gastrointest Liver Physiol 294, G610–618 (2008).

37. P. T. Pfluger, D. Herranz, S. Velasco-Miguel, M. Serrano, M. H. Tschop, Sirt1 protects against high-fat diet-induced metabolic damage. Proc Natl Acad Sci U S A 105, 9793–9798 (2008).

38. J. B. Weir, New methods for calculating metabolic rate with special reference to protein metabolism. J Physiol 109, 1–9 (1949).

39. C. Canto, P. M. Garcia-Roves, High-Resolution Respirometry for Mitochondrial Characterization of Ex Vivo Mouse Tissues. Curr Protoc Mouse Biol 5, 135–153 (2015).

40. G. Oemer et al., Molecular structural diversity of mitochondrial cardiolipins. Proc Natl Acad Sci U S A 115, 4158–4163 (2018).

41. T. Pluskal, S. Castillo, A. Villar-Briones, M. Oresic, MZmine 2: modular framework for processing, visualizing, and analyzing mass spectrometry-based molecular profile data. BMC Bioinformatics 11, 395 (2010).

42. C. W. Law, Y. Chen, W. Shi, G K. Smyth, voom: Precision weights unlock linear model analysis tools for RNA-seq read counts. Genome Biol 15, R29 (2014).

43. S. Falcon, R. Gentleman, Using GOstats to test gene lists for GO term association. Bioinformatics 23, 257–258 (2007).

44. F. Supek, M. Bosnjak, N. Skunca, T. Smuc, REVIGO summarizes and visualizes long lists of gene ontology terms. PLoS One 6, e21800 (2011).

